# Ingestion-activated CGRP neurons control learning but not satiety

**DOI:** 10.1101/2025.10.08.681275

**Authors:** Brooke C. Jarvie, Anagh Ravi, Rhea Nanavati, Ho Namkung, Longhui Qiu, Truong Ly, Jun Y. Oh, Olivia K. Barnhill, Zachary A. Knight

**Affiliations:** Department of Physiology, University of California, San Francisco, San Francisco, CA 94158, USA; Kavli Center for Fundamental Neuroscience, University of California, San Francisco, San Francisco, CA 94158, USA; Neuroscience Graduate Program, University of California, San Francisco, San Francisco, CA 94158, USA; Howard Hughes Medical Institute, University of California, San Francisco, San Francisco, CA 94158, USA

## Abstract

Calcitonin Gene-Related Peptide (CGRP) neurons in the parabrachial nucleus are critical for sickness and malaise but have also been proposed to control non-aversive (rewarding) satiation. How one cell type can coordinate these opposing processes has not been explained. Here we reinvestigate the function of these cells using single-cell imaging as well as optical and chemogenetic manipulations. Contrary to current models, we show that CGRP neurons do not track cumulative food consumption, and their activity is not necessary for meal termination.

Instead, we identify two distinct populations of CGRP cells, one of which responds rapidly to appetitive signals during ingestion and the other of which responds slowly to aversive visceral cues. Surprisingly, the ingestion-activated CGRP neurons are important for learning about post- ingestive effects but have little effect on food consumption. This reveals two populations of CGRP neurons that are sequentially engaged during, and responsible for, the distinct stages of post-ingestive learning.

## Introduction

The decision to stop eating can arise from either satiety, which is rewarding, or overconsumption and sickness, which are aversive. How these distinct internal states are differentially represented in the brain is a longstanding and unresolved question in neuroscience.

Calcitonin Gene-Related Peptide (CGRP) neurons in the parabrachial nucleus (PBN) are a cell type that sits at the interface between sickness and satiety. Stimulation of CGRP neurons robustly inhibits feeding, and these neurons are activated by an array of signals that induce nausea or visceral malaise^1–4^. Moreover, CGRP neuron stimulation is itself aversive, and activation of these cells is necessary and sufficient for learning about many aversive cues^4–10^. This has led to the hypothesis that CGRP neurons function as a “general alarm” that signals danger, especially as it relates to internal state^11^.

Nevertheless, a separate line of research has suggested that CGRP neurons also play an important role in non-aversive physiologic satiation and meal termination. This is supported by the finding that silencing CGRP neurons delays meal termination and can also attenuate behavioral responses to injection of gut hormones that promote satiety^3^. Consistent with this, one study performed single-cell imaging of CGRP neurons and found that virtually all these cells become activated between 30-50 minutes after the start of a meal^12^. However, the same study also reported that >90% of CGRP neurons are activated by diverse stimuli associated with pain and sickness, raising the question of how the same population of cells can represent both rewarding and aversive internal states.

Here, we have reinvestigated how CGRP neurons are regulated in different behavioral contexts in order understand how they control both satiety and sickness.

## Results

### A subset of CGRP neurons is rapidly activated in a manner time-locked to ingestion

To investigate the regulation of PBN-CGRP neurons during food consumption, we targeted expression of GCaMP6s to these cells by injecting a Cre-dependent AAV into the PBN of *Calca*^Cre:GFP^ mice^1^ and in the same surgery installed a GRIN lens for microendoscopic imaging (**Fig. 1a**, **Supplemental Fig. 1a**). We then gave hungry mice access to the complete liquid diet Ensure and performed single-cell calcium imaging during self-paced ingestion (30 minutes).

**Figure 1.**
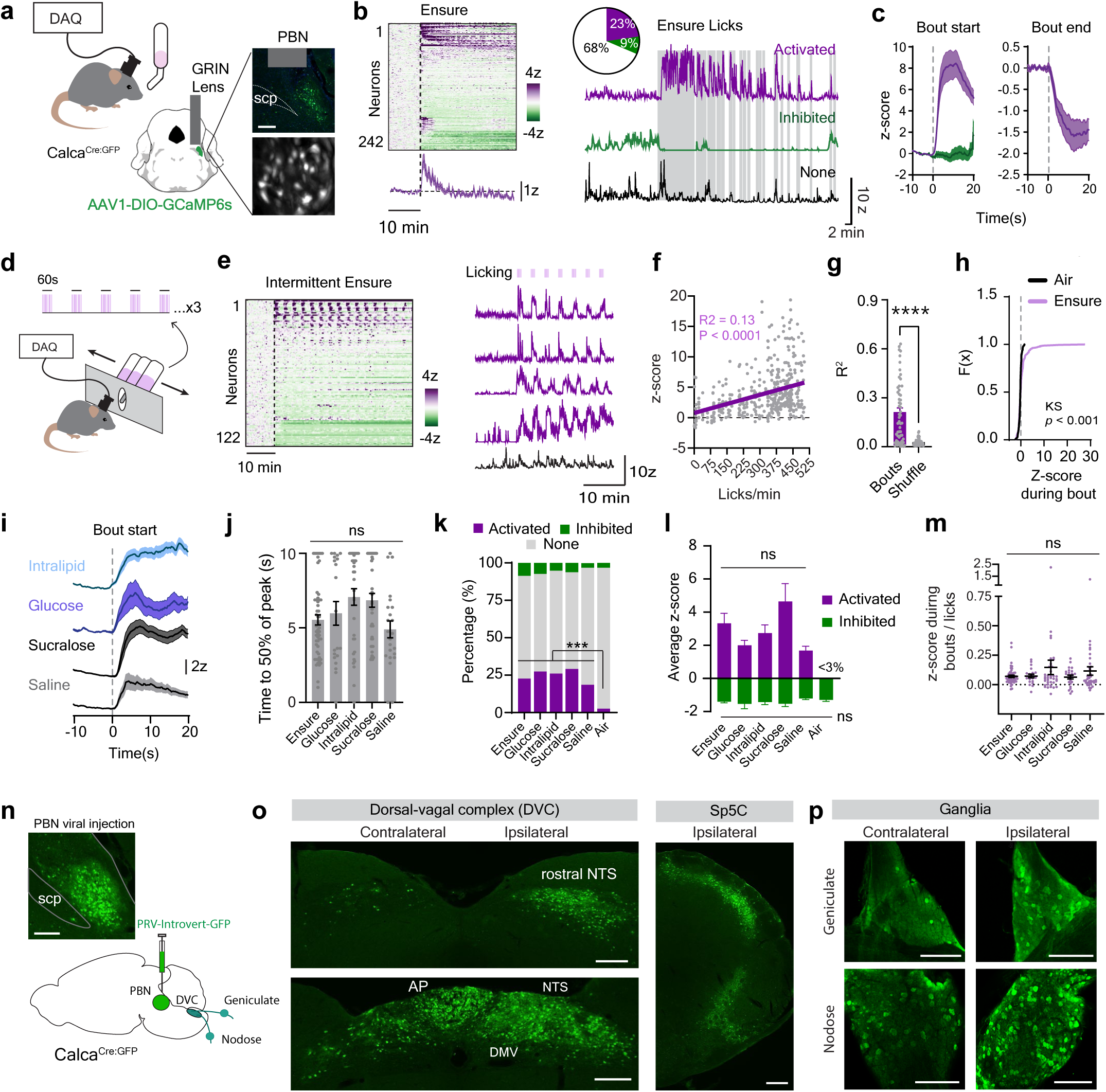
A subset of CGRP neurons tracks ingestion dynamics. **a**, Single-cell imaging of CGRP neurons in the parabrachial nucleus (PBN) during self-paced licking. Right, Representative lens placement and GCaMP6s expression in CGRP neurons and view through the GRIN lens. scp, superior cerebellar peduncle. **b**, Left, heatmap and average CGRP neuron population response to Ensure consumption (*N* = 10 mice). Right, quanitfication and example of individual neural responses to Ensure. Vertical gray bars represent licks. **c**, Peri-stimulus time histogram (PSTH) of CGRP neuron responses relative to the first and last lick in a bout. **d,** Schematic of mice being given intermittent access to Ensure on the Davis Rig. **e**, Heatmap of neural responses to intermittent licking and example neural traces (*N* = 5 mice). **f**, Correlation between licking and mean z-scored response. **g**, R^2^ of lick-bout state predicting neural activity of the activated population (observed vs. shuffle control). Each point represents one neuron. **h**, CDF of CGRP neuron responses during licking bouts for air vs Ensure. **i**, PSTH of activated neurons in response to licking for various solutions. **j**, Time to 50% of peak response following first lick in a bout. **k**, Quantification of neural responses. **l**, Average z-score during the first 10 minutes of solution access of activated and inhibited neurons. **m**, Average neural responses during bouts, normalized to the number of licks. **n**, Injection of the Cre-dependent multi-synaptic retrograde pseudorabies virus with GFP (PRV-Inrovert-GFP), and a representative image of viral expression in the PBN. **o**, Expression of PRV-Introvert-GFP in cells upstream of CGRP neurons in the dorsal vagal complex and the caudal spinal trigeminal nuclues (Sp5C) 3 days after injection of PRV-Introvert-GFP (*N* = 3 mice). **p**, PRV-Introvert-GFP expression in geniculate and nodose ganglia. Data are represented as mean ± SEM. AP, area postrema. NTS, nucleus of the solitary tract. DMV, dorsal motor vagus. Scale bars, 200 µm. ns means not significant, *** *p* < .001, **** *p* < .0001. One-way ANOVA (j, l, m) and Fisher exact tests with bonferroni correction (k).

We found that 23% (55/242 neurons from 10 mice) of CGRP neurons were rapidly activated during Ensure consumption in a manner that was time-locked to licking (mean bout activation: 3.3 ± 0.6 z, **Fig. 1b-c**). This lick-triggered activation began with the first lick, ramped over several seconds (50% of peak activation at 5.5 ± 0.3 s) and then gradually declined when the lick bout ended (50% decrease after 6.7 ± 0.4 s, **Fig. 1c**). Similar time-locked, lick-triggered responses were observed when mice were given intermittent access to Ensure in one-minute trials (**Fig. 1 d-e**) and the magnitude of this response scaled with the size of the lick bout (R^2^ = 0.130, *p* < 0.0001, **Fig. 1f**). These responses were consistent at the single-cell level, with activated neurons responding in 79 ± 3% of licking trials. Moreover, much of the variance in single neuron activity could be explained by whether the animals were licking (R^2^ = 0.27 ± 0.02 vs 0.04 ± .003, bouts vs. shuffle, *p* < 0.0001, **Fig. 1g**). Thus, many CGRP neurons are activated during Ensure ingestion in a manner time-locked to licking.

We next sought to define the sensory signals that activate CGRP neurons during ingestion. First, we noted that CGRP neuron activation was not simply due to the mechanical act of licking, because less than 3% of neurons responded when mice performed “air licking” at an empty bottle (KS test of CGRP neuron response during bouts of licking for air vs. Ensure, *p* = 0.0002, **Fig. 1h**). This response was also not macronutrient-specific, because consumption of a pure fat solution (Intralipid) or a pure sugar solution (glucose) caused time-locked activation of CGRP neurons (**Fig. 1i-j**, **Supplemental Fig. 1b**) that was similar to Ensure in both the percentage of responsive cells (**Fig. 1k**) and the strength of their activation (mean z-score: Ensure 3.3 ± 0.6, Intralipid 2.7±0.5, glucose 2.0±0.3, comparison of all conditions *p* = 0.55, **Fig. 1l**). We also observed strong CGRP neuron activation during consumption of the non-nutritive sweetener sucralose and ingestion of saline (a mildly salty solution, **Fig. 1i-l**). Mice sampled all solutions provided, and, despite some differences in volume consumed (**Supplemental Fig. 1c**), normalizing the neural activation during ingestion to the number of licks did not reveal any differences between the solutions tested (**Fig. 1m**). We also observed a small subset of CGRP neurons that were tonically inhibited during our imaging sessions, but these responses did not correlate with ingestion (**Fig. 1k-l**, **Supplemental Fig. 1b**). Taken together, these data indicate that a subset of CGRP neurons are rapidly activated during ingestion by a signal that depends on the volume consumed but is independent of nutrient identity or caloric value.

### CGRP neurons receive extensive input from pregastric sensory pathways

The time-locked activation of CGRP neurons during ingestion suggests that these neurons may be modulated by sensory input from the mouth or esophagus^13–16^. To identify possible sources of this pregastric feedback, we performed polysynaptic retrograde tracing to visualize the inputs to CGRP neurons. We did this by injecting a Cre-dependent pseudorabies virus expressing GFP (PRV-Introvert-GFP, **Fig. 1n**)^17^ into the PBN of *Calca*^Cre:GFP^ mice and examining the spread of virus at timepoints 2-4 days after infection (**Fig. 1o-p**, **Supplemental Fig. 2a-b**).

On day 2, there was extensive labeling of neurons in regions associated with visceral malaise (area postrema)^18–20^, taste (rostral nucleus of the solitary tract (NTS))^21–23^, and feedback from the esophagus and lower gastrointestinal tract (medial and caudal NTS^24,25^, **Supplemental Fig. 2b**)^9,26–29^. On days 3 and 4, we observed the appearance of GFP expression in peripheral sensory neurons. This included labelling in the geniculate ganglia, which contains the cell bodies of the neurons that carry gustatory information from the facial nerve (**Fig. 1p**)^30^ and the nodose ganglia, which contains the cell bodies for afferent vagal neurons that innervate the esophagus and lower gastrointestinal tract^24,31^. We also observed strong labelling in the caudal part of the spinal nucleus of the trigeminal nerve, which receives pain and temperature information from the oral cavity and face^32^. For all these structures, labeling was stronger on the side ipsilateral to the injection site (**Fig. 2o-p**) and was specific to inputs to CGRP neurons, as no GFP expression was observed after injection of PRV-Introvert-GFP into wild-type mice (**Supplemental Fig. 2c**). Thus, CGRP neurons receive prominent input from sensory pathways that innervate the oral cavity and esophagus as well as the gastrointestinal tract, consistent with the observation that a subset of these neurons is rapidly activated during ingestion.

**Figure 2.**
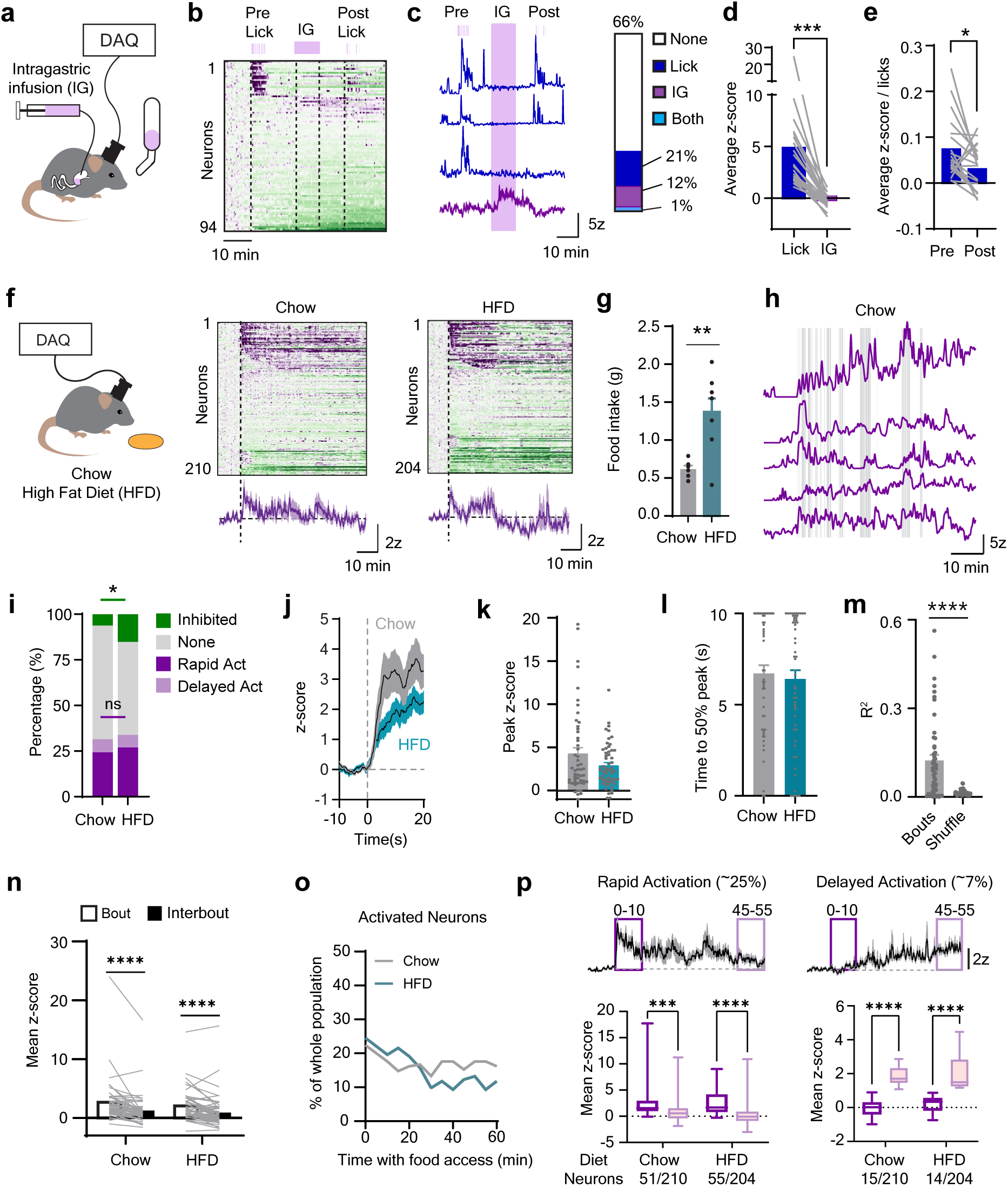
CGRP neurons do not track cumulative food intake. **a**, Mice licked for Ensure before and after a 1 mL intragastric infusions of Ensure during single-cell calcium imaging of CGRP neurons. **b**, Heatmap showing neural responses to either licking or IG infusion (*N* = 6 mice). **c**, Example traces and quantification of neural responses. **d**, Average z-score during either licking or IG infusion for ingestion-activated neurons. **e**, Mean lick-normalized responses to Ensure pre vs. post IG infusion. **f,** Left, Schematic and right, heatmaps and average CGRP neuron population response to intake of chow (*N* = 6 mice) or high fat diet (HFD, *N* = 7 mice). **g**, Amount of food consumed during the imaging session. **h**, Example responses of CGRP neurons to food intake. Vertical gray bars represent bites. **i**, Quanitfication of neural responses to food ingestion. **j**, PSTH average of CGRP neuron responses relative to the start of a bout. **k**, Peak z-score of the PSTH for each neuron. **l**, Time to 50% of the peak response following the start of a bout. **m**, Variance in single-cell activity explained by bout state across activated neurons. **n**, Comparison of z responses during and between bouts. **o**, Percentage of neurons that were activated in 5-min bins over time. **p**, Average neural traces and the mean z response during the first and last 10 minutes epochs of food access of neurons that were activated when first given food access (rapid) or at the end of the session (delayed). ns means not significant, * *p* < 0.05, ** *p* < 0.01, *** *p* < .001, **** *p* < .0001.

### Activation of CGRP neurons during licking is not potentiated by post-ingestive signals

If CGRP neurons drive satiation^3^, then their activity would be predicted to increase as the meal progresses. One way this could happen is that post-ingestive signals potentiate CGRP neuron responses to licks, such that this time-locked activation increases in magnitude as the meal proceeds (**Supplemental Fig. 1d**). However, we observed little evidence for this during self-paced ingestion. For example, lick responses for Ensure during intermittent access tests were similar in magnitude at the beginning and end of the 60 min session (0.015 ± 0.003 vs. 0.013 ± 0.004 mean z-score per lick, *p* = 0.56, **Supplemental Fig. 1e**), despite the fact that mice consumed a large volume of Ensure that would be predicted to promote satiation (5237 ± 537 licks, *N* = 4 mice).

To test this idea more rigorously, we equipped mice with intragastric (IG) catheters^33,34^ so that we could infuse nutrients directly into the stomach while recording CGRP neuron dynamics by microendoscopy (**Fig. 2a**). We then measured how lick responses were altered by IG infusion of nutrients. To do this, hungry mice were first given brief access (5 min) to Ensure for oral consumption; the lick bottle was then removed and the mice were given an IG infusion of Ensure (1 mL over 10 min); the lick bottle was then returned and mice were given a second brief access period (5 min) for Ensure oral consumption (**Fig. 2b-d**).

We found that mice consumed more Ensure in the first access period than in the second (1096 ± 83 vs. 478 ± 111 licks, session 1 vs. 2, *p* = 0.002), and this was due to a decrease in the number of lick bouts initiated (12 ± 2 vs. 7 ± 2 bouts, session 1 vs. 2, *p* = 0.03). This confirms that the IG infusion was sufficient to promote satiation. However, the activation of CGRP neurons during licking was reduced, rather than increased, following IG nutrient infusion. This was due to both a decrease in the number of Ensure-responsive neurons (21% vs. 14%, from 6 mice, **Fig. 2c**) and a decrease in the activation of those neurons per lick (0.07 ± 0.01 vs 0.03 ± 0.01 mean z-score per lick, *p* = 0.01, **Fig. 2e**). Thus, post-ingestive nutrients do not potentiate the rapid activation of CGRP neurons during licking, as might be expected if this activation functions to promote satiation.

### The tonic activity of CGRP neurons does not track cumulative food intake

An alternative mechanism by which CGRP neurons could promote satiation is by tracking cumulative food intake in their tonic or baseline activity. This tonic activation would be detected as an increase, as the session progresses, in the activity of CGRP neurons between bouts of licking (**Supplemental Fig. 1f**). However, we found little evidence for this (**Supplemental Fig. 1g-j**). For example, during intermittent access tests, we only detected one CGRP neuron (out of 122 cells) that appeared to show ramping activation during ingestion (shown in **Fig. 1e**).

The preceding experiments all used liquid diets, but it is possible that CGRP neurons respond differently to solid food consumption. We therefore allowed fasted mice to consume either normal chow or high-fat diet (60% HFD) and imaged CGRP neuron responses (**Fig. 2f-h**). We found that ∼25% of neurons were rapidly activated during consumption of either chow or HFD (chow: 51/210 neurons from 6 mice, HFD: 55/204 neurons from 7 mice, **Fig. 2i**). Similar to what we observed with liquid diets, most CGRP neuron activation during solid food consumption was time-locked to bouts of ingestion (**Fig. 2j-n**). Moreover, there were no differences between chow and HFD in this bout-triggered activation (**Fig. 2j-n**).

It was previously reported that almost all CGRP neurons become progressively activated between 30-50 minutes after the start of food consumption^12^. However, we observed no widespread activation on this timescale, even when mice consumed a large quantity of HFD (1.37 ± 0.50 g, **Fig. 2g, o**). Rather, we found that the mean activity of ingestion-responsive neurons actually decreased as the trial progressed (2.3 ± 0.36 v 0.85 ± 0.36 z, first v. last 10 minutes, *p* < 0.001 ,**Fig. 3p**). We did identify a small subset of neurons that were not initially responsive to ingestion but became more active as mice consumed chow or HFD (7% of neurons exhibiting delayed activation to both diets, **Fig. 2i, p**). However, a larger percentage of neurons became inhibited with continued food intake (6.6 ± 2.7 to 30.5 ± 5.9% by the end of the session, *p* = 0.007, **Supplemental Fig. 3a-d**). Thus, the baseline activity of most CGRP neurons does not increase with total food consumed, suggesting that their activity is not driven by post-ingestive signals during a normal meal.

**Figure 3.**
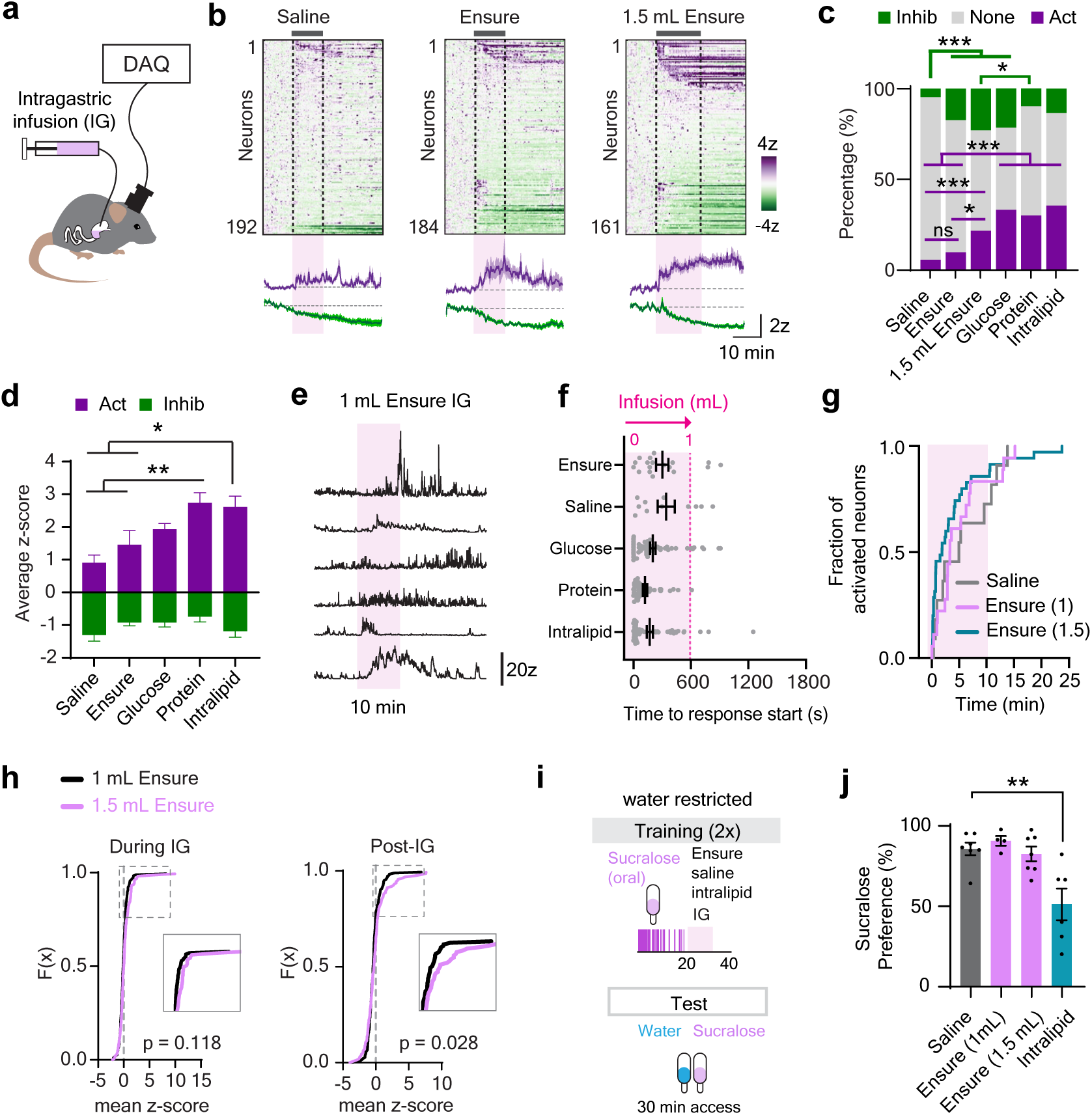
CGRP neurons show limited responses to post-ingestive nutrients. **a**, Single-cell imaging of CGRP neural responses to intragastric (IG) infusions. **b**, Heatmaps of neural responses to 1.0 mL saline (N = 7 mice), 1.0 ((N = 5 mice) or 1.5 mL Ensure IG infusions (N = 7 mice). **c**, Quantification of neural responses to IG infusions. **d**, Mean z-score of neural responses during IG infusions of IG-activated and inhibited neurons. **e**, Example neural traces in response to a 1.0 mL Ensure IG infusion. **f**, Time to the response onset relative to the start of the IG infusion. **g**, Cumulative fraction of response times of activated neurons. Log-rank *p* = 0.557. **h**, Distribution of mean z responses during the first 10 minutes of the IG infusion or 10 minutes post infusion. **i**, Paradigm for conditioned taste avoidance where sucralose is paired with an IG infusion of different solutions. **j**, Decreased preference for sucralose in a two-bottle choice test after pairings with IG intralipid, but not other solutions. **** *p* < .0001, *** *p* < .001, ** *p* < .01. * *p* < .05.

### CGRP neurons respond to IG nutrient infusions under conditions that cause malaise

These findings are contrary to the conventional wisdom that CGRP neurons track gastrointestinal signals, such as stomach distension, to trigger meal termination^1,3,35^. We therefore tested whether CGRP neurons receive gastrointestinal signals directly by measuring responses of CGRP neurons to infusions of different solutions into the stomach.

We first recorded neural responses to IG infusion of Ensure, which we delivered in an amount (0.93 kcal; 0.1 mL/min over 10 min) that approximates a moderately-sized meal and has been shown to modulate many neural cell types that control food intake^13,14,24^. We found that only a small percentage of CGRP neurons (18/184 neurons from 5 mice) were activated by Ensure IG infusion (**Fig. 3a-g**), and this was not different from the response to equivolemic saline infusion (10% vs. 6% activated neurons (11/192), Ensure vs. saline, *p* = 1, **Fig. 3c**). Increasing the number of calories infused, either by increasing the volume (1.5 mL Ensure, 1.40 kcal) or by infusing pure fat (1.0 mL Intralipid, 2 kcal), resulted in activation of a greater percentage of CGRP neurons (**Fig. 3c, h**, **Supplemental Fig 4a**). We also observed, unexpectedly, that infusing pure solutions of sugar (glucose, 24%) or protein (ProteinEx, 45%) induced stronger CGRP neuron activation than infusions of calorie and volume-matched solutions of a complete diet (**Fig. 3c-d**, **Supplemental Fig. 4a-e**). This indicates that CGRP neurons can respond to IG infusions when delivered in large amounts or when there are nutrient imbalances. Nevertheless, this response to IG infusion appeared to be specific to the IG infusion paradigm, as we did not observe sustained activation of CGRP neurons when the same solutions were consumed by mouth in comparable or greater quantities (**Fig. 1i-m**, **Supplemental Fig. 1b-c**). Consistently, the CGRP neurons activated by oral consumption were different from the CGRP neurons activated by IG infusion (**Fig. 2b-d**). Thus, the activation of CGRP neurons by IG infusion does not seem to mimic normal ingestion.

Given that CGRP neurons are associated with visceral malaise^1,2,4,6^, we reasoned that IG infusions that activate CGRP neurons might be aversive. To test this, we measured whether IG infusion of different solutions causes a conditioned taste avoidance (CTA). Mice were trained by pairing, on two separate days, a novel tastant (flavored sucralose) with subsequent IG infusion of either saline (control), Ensure (1.0 or 1.5 mL), or Intralipid (1.0 mL, **Fig. 3i**). Mice then underwent a two-bottle preference test for sucralose and water. This revealed that IG infusion of Intralipid, which strongly activates CGRP neurons, reduced the preference for paired sucralose (85.7% ±4.0 vs. 51.3 ±9.9% preference, IG saline vs. Intralipid, *p* = 0.002), whereas there was no effect of IG infusions of Ensure (**Fig. 3j**). In a separate experiment, we found that injection of the GLP1R agonist Exendin-4 induced substantial activation of CGRP neurons only at higher doses that become more aversive (**Supplemental Fig. 5a-d**)^36–38^. Taken together, these data suggests that the ability of CGRP neurons to respond to certain post-ingestive signals is more related to malaise than satiation.

We wondered whether the stronger activation of CGRP neurons by IG Intralipid might be due to differences in gastric emptying. To test this, we gave mice an IG infusion of either saline, Ensure or Intralipid (1.0 mL over 10 min) and then measured the contents in different parts of the GI tract (**Supplemental Fig. 6a-c**). As expected, Ensure and Intralipid emptied from the stomach slower than saline **(Supplemental Fig. 6d**), and, correspondingly, stomach size and the weight of the remaining stomach contents were greater for these solutions compared to saline (**Supplemental Fig. 6b,e**). However, there was no difference in any of these parameters between Ensure and Intralipid. There was also no difference in the amount of Ensure versus Intralipid that reached the duodenum or jejunum during the experiment (**Supplemental Fig. 6b**). Thus, differences in GI motility are unlikely to explain the differential response of CGRP neurons to these solutions.

### Different populations of CGRP neurons respond to ingestion and visceral malaise

The discovery that a subset of CGRP neurons is rapidly activated by appetitive signals associated with ingestion (**Fig. 1**) raises the question of whether these are the same cells that respond to aversive signals associated with sickness. To test this, we gave fasted mice access to Ensure (10 min) followed by an i.p. injection of LiCl (84 mg/kg), which is commonly used to induce visceral malaise^4,6,39^ and activates CGRP neurons^1,10^. We found that a large subpopulation of CGRP neurons were activated by LiCl (42%, 83/196 neurons from 8 mice, 4.5 ± 0.49 z, **Fig. 4 a-c**), and these cells were largely non-overlapping with the neurons activated by Ensure (only 5% of neurons responding to both stimuli). To confirm that aversive and appetitive signals activate different CGRP neurons, we repeated this experiment using Ensure followed by injection of “vomitoxin” (deoxynivalenol, or DON), a mycotoxin that causes gastrointestinal distress and nausea (**Fig. 4d-f**)^40^. We found that 33% (36/108 neurons from 5 mice) of CGRP neurons were activated by DON (3.9 ± 0.48 average z), and these DON activated neurons were mostly non-overlapping with the cells that responded to Ensure (only 3% responding to both, 3/108 neurons). Thus, aversive and ingestive signals activate different subsets of CGRP neurons in the PBN.

**Figure 4.**
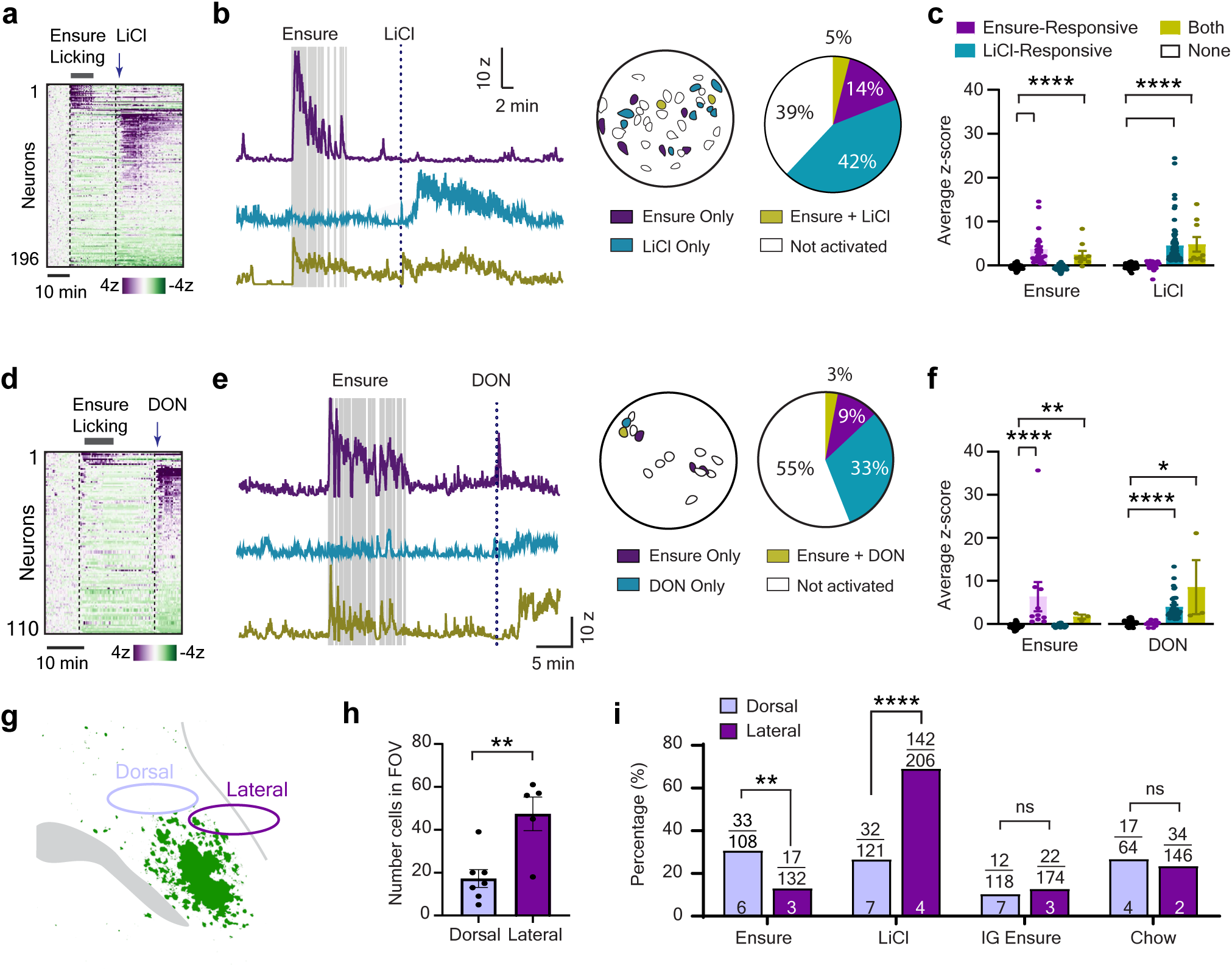
Distinct CGRP neurons respond to ingestion and malaise. **a**, Heatmap of neural responses to self-paced licking for Ensure and subsequent LiCl administration in the same recording session (*N* = 8 mice). **b**, Left, example responses of neurons that responded to Ensure, LiCl, or both stimuli. Vertical gray bars are licks. Right, spatial distribution of cells with each response type in one animal, and quantification of the percentage of each type of response across all mice. **c**, Quantification of the average z-score during different stimuli, for each category of neuron. **d**, Heatmap of neural responses to Ensure and DON (*N* = 5 mice). **e**, Example neural traces and quantification of responses. **f**, Mean z-score response of neurons based on the stimuli they respond to. **g,** Schematic showing the two regions where GRIN lenses were placed for recordings. **h**, Quantification of the number of neurons per animal based on the location of the lens. **i**, The percentage of neurons responding to different stimuli based on the lens location. The ratio is responsive cells divided by total cells, from the number of animals indicated by the numbers in the bars. Analysis is a Fisher’s Exact Test for each stimuli. **** *p* < .0001, *** *p* < .001, ** *p* < .01, * *p* < .05.

We wondered whether the CGRP neurons that respond to ingestion and malaise are spatially segregated in the PBN. Post-hoc analysis of our GRIN lens placements revealed that they were clustered into two groups, one located more dorsally and one more laterally within the PBN (**Supplemental Fig. 1a**, **Fig 4g-h**). Reanalysis of neural responses based on lens placement revealed that more neurons responded to LiCl in the lateral region (69%, 142/206 neurons from 4 mice) compared to the dorsal region (26%, 32/121 neurons from 7 mice, Fisher Exact Test, *p* < 0.0001, **Fig. 4i**). On the other hand, more neurons were activated by Ensure consumption in the dorsal region compared to the lateral (33/108 cells from 6 mice vs. lateral recordings with 17/132 neurons from 3 mice, *p* <0.01). However, we did not see any differences between these two regions in response to intake of solid food or 1.5 mL IG Ensure infusions (**Fig 4i**). These data indicate that there is some topographic organization of CGRP neuron responses in the PBN, but this is not absolute and does not apply to all stimuli.

### The activation of CGRP neurons during ingestion is dispensable for satiation

We next sought to determine the function of the ingestion-triggered activation of CGRP neurons. To do this, we used two complementary approaches to silence CGRP neurons while mice consumed food (optogenetics using GtACR and chemogenetics using hM4D; **Fig. 5a-f**, **Supplemental Fig. 7a-d** and **Supplemental Fig. 8a-j**). We performed these silencing experiments under a variety of conditions, including in both fasted and fed mice and with both solid and liquid food. We also measured the effect of closed-loop (lick-triggered) optogenetic silencing, which blocks the ingestion-triggered bursts in CGRP neuron activity, and compared this to tonic inhibition. We analyzed these data to look for effects on total food consumed, meal patterns (meal duration and number) and bout microstructure (bout duration and number).

**Figure 5.**
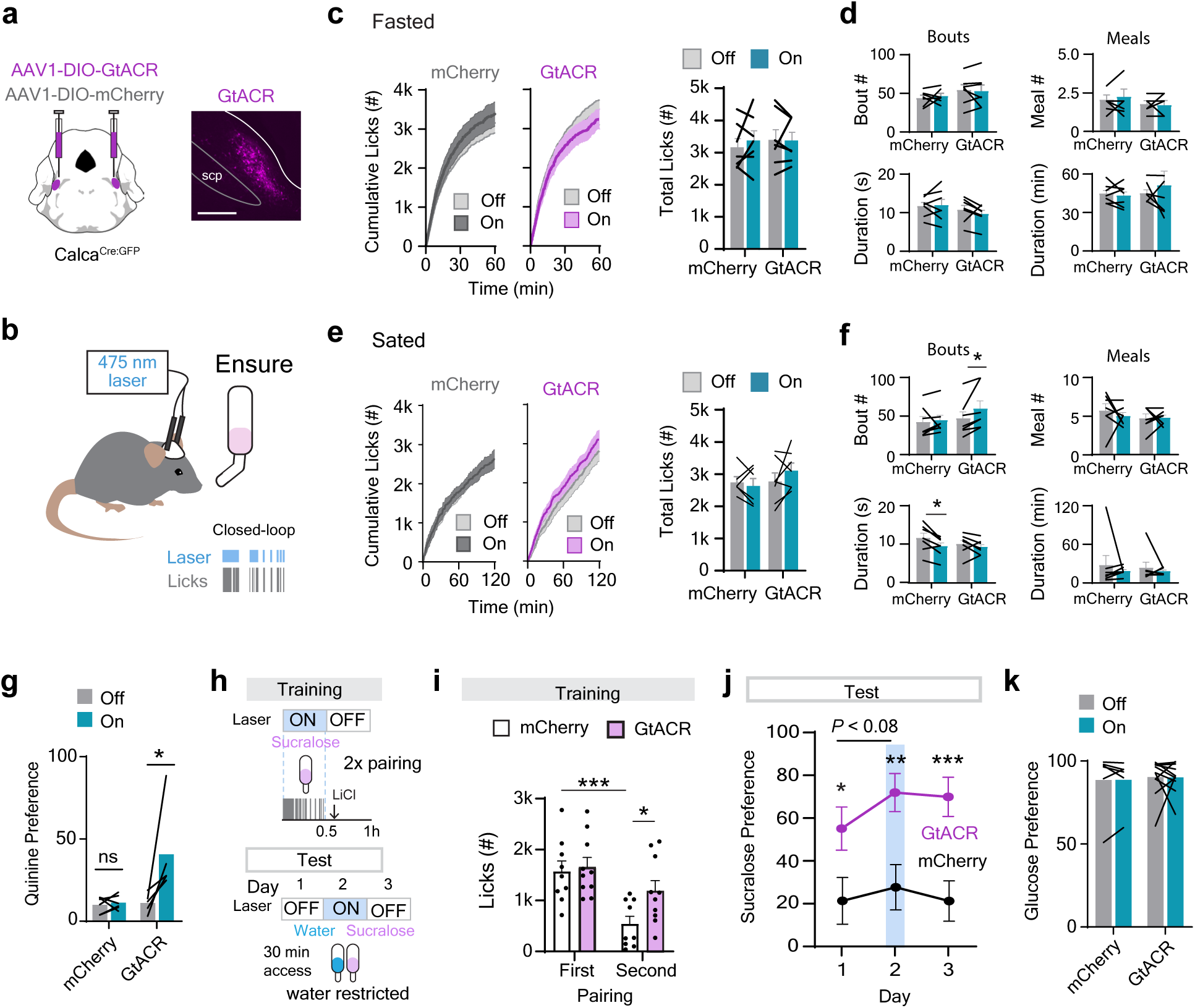
Rapid ingestion responses are important for learning but not satiation. **a**, *Calca*^Cre^ mice were injected with AAV1-DIO-GtACR or AAV1-DIO-mCherry and implanted with optic fibers. Right, image of GtACR expression in the PBN, scale bar 200 um. **b**, Schematic of mice receiving lick-triggered closed-loop optogenetic inhibition while given access to Ensure. Vertical gray bars are licks. **c**, There were no differences in total number of licks for Ensure in fasted mice. **d**, There were no changes in bout or meal number and duration in fasted mice. **e**, There were no differences in cumulative rate or total lick number in sated mice given Ensure access. **f**, CGRP silencing increased bout number but not bout duration, and meal number and duration were not different between groups. **g**, Continuous GtACR-inhibition of CGRP neurons increased preference for a bitter solution vs. water. **h**, Training and testing paradigm for CTA where CGRP neurons were selectively inhibited during sucralose access during training. **i**, GtACR mice drank more sucralose than control mCherry mice on the second day of training. **j**, GtACR mice showed greater sucralose preference in two-bottle choice tests across consecutive days, where CGRP neurons were inhibited on day 2. **k**, CGRP inhibition had no effect on sweet preference in a two-bottle choice test. scp, superior cerebellar peduncle. * *p* < .05, \*\**p* < .01, \*\*\**p* < .001.

The overall result from these experiments was that CGRP neuron silencing had little or no effect on food consumption under most conditions tested (summarized in **Table 1**). Specifically, in seven out of eight experiments, CGRP neuron silencing had no effect on the total amount of food consumed. In one experiment (consumption of Ensure by fed mice over two hours), there was a small increase in food intake following chemogenetic silencing with hM4D (2212± 176 vs 2680 ± 263 licks saline vs. DCZ within-subject, *p* = 0.02), which was associated with an increase in bout number (**Supplemental Fig. 8b-c**). However, we did not observe an increase in food intake under the same conditions when we silenced using a different method (GtACR, either closed-loop or tonic inhibition; **Fig. 5e-f**, **Supplemental Fig. 7c-d**). In no experiment did we observe a change in meal duration, which was the parameter previously reported to be under the control of CGRP neurons^3^. These negative results were not due to inefficient silencing, because we confirmed the efficacy of silencing for both methods using positive controls. Specifically, we showed that silencing CGRP neurons using optogenetics increased consumption of a bitter quinine solution (**Fig. 5g**)^9,41^, whereas silencing CGRP neurons using chemogenetics blocked CTA induced by LiCl injection (**Supplemental Fig. 8i-j**)^4,6^. These data indicate that CGRP neurons are not required for satiation and play little or no role in the control of food intake, at least under the conditions tested here.

**Table 1.** Summary of effects of CGRP neuron silencing on food intake. Comparison of food intake and meal microstructure between experimental mice (GtACR or hM4D- expressing) and control mice expressing mCherry in CGRP neurons in the PBN. ↑, increase; =, no differences between groups or conditions. N/D, no data.

| Food | State | Tool | Inhibition | Total Intake | Bout # | Bout duration | Meal # | Meal duration |
| --- | --- | --- | --- | --- | --- | --- | --- | --- |
| Ensure | Fasted | GtACR | Closed-loop | = | = | = | = | = |
|  | Fasted | GtACR | Continuous | = | = | = | = | = |
|  | Fasted | hM4Di | Continuous | = | = | = | = | = |
|  | Fed | GtACR | Closed-loop | = | ↑ | = | = | = |
|  | Fed | GtACR | Continuous | = | = | = | = | = |
|  | Fed | hM4Di | Continuous | ↑ | ↑ | = | = | = |
| Solid Food | Fasted | hM4Di | Continuous | = | N/D | N/D | N/D | N/D |
|  | Baseline | hM4Di | Continuous | = | N/D | N/D | = | = |

### CGRP neuron activity during consumption is required for post-ingestive learning

We next considered the possibility that the ingestion-triggered activation of CGRP neurons is involved in learning. In this regard, CGRP neurons are already known to be critical for CTA^4,6^ (**Supplemental Fig. 8j**), and it is generally assumed that this is because CGRP neurons supply the unconditioned stimulus (US; e.g. LiCl) in this paradigm. Consistently, CGRP neurons are activated by LiCl^1,10^ (**Fig. 4a-c**) and their artificial activation can substitute for LiCl in driving CTA^4,6,10^. On the other hand, the conditioned stimulus (CS, e.g. taste) in this paradigm is thought to be encoded by parallel and downstream brain regions^10,39,42^. However, the discovery that a subset of CGRP neurons is activated by appetitive tastants (**Fig. 1**) led us to hypothesize that CGRP neurons might also supply information about the CS during learning.

To test this hypothesis, we took advantage of the fact that optogenetic silencing enables inhibition of CGRP neurons selectively during exposure to the CS but not the US. Mice expressing GtACR in CGRP neurons (or mCherry for controls) were trained by exposure to a novel tastant (sucralose, 30 min) followed by an injection of LiCl, repeated across two training sessions. Both groups received laser stimulation during sucralose exposure, but not LiCl treatment (**Fig. 5h**). Remarkably, we found that inhibiting CGRP neurons only during sucralose consumption was sufficient to block the development of CTA, as reflected by increased consumption of sucralose on day 2 of training (**Fig. 5i**) and increased preference for sucralose in a subsequent two-bottle choice test (**Fig. 5j**, day 1). Inhibition of CGRP neurons during a later two-bottle choice test further increased preference for sucralose, in agreement with the previously reported role of CGRP neurons in the retrieval of a CTA memory (**Fig. 5j**, day 2)^4^.

Importantly, CGRP neuron inhibition did not interfere with the valence of sweet taste itself, as there was no difference in sucralose intake on the first day of training (1567 ± 210 vs. 1658 ± 186 licks, control vs. GtACR, *p* = 0.74; **Fig. 5i**) nor did it have an effect on preference for an alternate sweet solution (**Fig. 5k**)^41^. Together with our imaging results (**Fig. 4**), these findings reveal that separate populations of CGRP neurons track the CS and US during aversive learning, and that making this association requires CGRP neuron activity not only during exposure to the aversive US but also during the appetitive CS (**Fig. 1**).

## Discussion

Satiation and sickness are closely connected interoceptive states. Overeating can quickly transform satiation into discomfort, and nausea is one of the major side effects of appetite- curbing drugs used to treat obesity. Thus, understanding how these two states are differentially represented in the brain is a longstanding and important question in physiology.

Parabrachial CGRP neurons are unique in that they are broadly implicated in the control of nausea and sickness^1,4,6^ but are also thought to contribute to normal satiation^3,12^. To understand how these two functions are performed by the same cells, we reinvestigated the dynamics and function of CGRP neurons during both feeding and malaise. Contrary to current models^35^, we find that CGRP neurons do not track cumulative food intake, and their activity is not necessary for meal termination or most aspects of self-paced food consumption. However, a subset of CGRP neurons is rapidly activated during feeding in a manner time locked to ingestion. We show that these ingestion-activated cells are distinct from the neurons that respond to visceral malaise, and their activation during CS consumption is required for animals to learn to associate a CS with aversive post-ingestive stimuli. These findings reveal a role for different subsets of CGRP neurons across multiple stages of aversive learning.

### CGRP neurons are dispensable for normal satiation

While many lines of evidence implicate CGRP neurons in the control of sickness, malaise and danger, their role in meal termination rests primarily on two observations. First, it was shown that nearly all CGRP neurons become activated from 30-50 minutes after the start of feeding^12^, which correlates roughly with the onset of post-prandial satiety. Second, it was shown that silencing of CGRP neurons with tetanus toxin increases meal size in mice that are hungry^3^.

These observations have led to the idea that CGRP neurons track post-ingestive feedback from the GI tract to regulate the timing of meal termination^35^.

We were unable to find evidence to support this model. Regarding neural dynamics, we found that CGRP neurons respond heterogeneously to every stimulus tested and that very few cells track cumulative food intake in the way previously reported^12^. Specifically, a subset of CGRP neurons (∼25%) were activated by ingestion of either liquid or solid food, but this response was transient and time-locked to eating, suggesting it reflects pre-gastric signals such as taste or esophageal stretch^14–16^ or, possibly, the initial arrival of food to the stomach^43^. Few neurons showed ramping activation consistent with post-ingestive feedback, and, consistent with this, IG nutrient infusions strongly activated CGRP neurons only under conditions that caused CTA. On the other hand, we found that aversive stimuli, such as LiCl and DON, activated a subset of CGRP neurons (33-42%) that was distinct from those that responded to ingestion. This heterogeneity is consistent with other data showing, for example, that ∼30% of CGRP neurons respond to quinine or fox urine, 45-50% respond to a looming stimulus or an unexpected sound, and ∼60% respond to a foot shock^9^. In contrast, the study that reported CGRP neurons track cumulative food intake^12^ found that these neurons respond homogeneously (>90% activated) to not only feeding but also a wide range of aversive stimuli.

To test the necessity of CGRP neurons in regulating food intake, we silenced these neurons using a combination of optogenetics and chemogenetics and then measured the effect on self-paced food consumption. We observed no effect of silencing on the amount of food consumed in 7/8 experiments performed, with one condition (Ensured consumption in fed mice) showing a small increase in total consumption following chemogenetic but not optogenetic inhibition.

Importantly, CGRP neuron inhibition did not alter meal size, which is the parameter that was proposed to require CGRP neuron activity^3^, in any of our experiments. We cannot explain why our results differ from a previous study using tetanus toxin to silence these neurons, but it is conceivable that chronic loss of CGRP neurons results in circuit-level adaptations that are important for feeding behavior. Alternatively, it is possible that tetanus toxin silencing has off-target effects that arise from its high efficiency^44^ combined with the low level of leaky expression from most Cre-dependent AAVs^45^. Of note, the only other study that measured the effect of CGRP neuron silencing on food intake used chemogenetics (hM4D) and, similar to our results, reported no change in consumption^1^. Taken together, these results show that CGRP neurons play a minimal role in regulating food consumption under conditions commonly used in the lab and suggest that satiation and malaise involve different neural populations^46^.

### CGRP neurons are required during both the CS and US in aversive learning

Extensive evidence implicates CGRP neurons in aversive learning. For example, it has been shown that CGRP neurons are activated by diverse stimuli associated with danger or sickness^1,6,9,11,12^, and we confirmed in our imaging preparation that a subset of CGRP neurons (33-42%) are strongly activated by emetics such as LiCl and DON (**Fig. 4**). CGRP neurons were also activated by IG Intralipid under conditions that induced a mild CTA (**Fig. 3**)^47^. These observations, together with data from functional manipulations^4,6^, are consistent with the conclusion that CGRP neurons encode the US in aversive post-ingestive learning.

In this context, we were surprised to find that a separate population of CGRP neurons was activated by ingestion of appetitive solutions, which typically function as the CS in models of post-ingestive learning. Moreover, blocking this activation by silencing CGRP neurons only during CS consumption prevented the formation of a CTA (**Fig. 5**). This indicates that the sequential activation of two separate populations of CGRP neurons – one that tracks the CS and one that tracks the US – is required for forming a taste memory.

We do not know how CGRP neuron activation during CS consumption contributes to CTA, but our data argue against a few potential mechanisms. First, silencing CGRP neurons had no effect on consumption of the CS during the first day of training, indicating that this learning defect is not indirectly due to a change in behavior. Second, we found that CGRP neurons were reliably activated by ingestion of familiar liquid diets that the mice had consumed many times, indicating that this activation does not encode primarily neophobia^12,48^ or novelty^10^. This is relevant because novel CS solutions tend to induce a stronger CTA due to the absence of latent inhibition^6,49,50^. Third, while it is possible that CGRP neurons encode some sensory features of the CS, our data suggest these neurons respond primarily to ingestion itself rather than specific gustatory cues (**Fig. 1**). Of note, CGRP neurons are also important for other forms of aversive learning, such as threat memory^5,9,51^, but CGRP neurons do not respond to an auditory CS prior to threat conditioning^9,12^. This suggests that the engagement of CGRP neurons during the CS is specific to ingestive learning.

We think it is most likely that CGRP neuron activation during CS consumption acts as a permissive signal that is required, in some way, for subsequent creation of the taste memory in downstream circuits. In this regard, it has recently been shown that activation of CGRP neurons during CTA training is sufficient to reactivate the representation of the recently consumed taste in the downstream central amygdala (CeA)^10^. Thus, it is possible that the activation of CGRP neurons during CS consumption is necessary to tag these downstream CeA neurons for subsequent reactivation. Testing this idea will require identifying features such as gene expression^52,53^, connectivity^54,55^ or other properties^56^ that distinguish the ingestion-activated CGRP neurons from other cells and enable targeted exploration of their function.

## Supporting information

Supplemental Statistics Table

## Acknowledgements

This work was supported by R01-DK106399, R01-DK145100, and R01-DK138127. Z.A.K. is an Investigator of the Howard Hughes Medical Institute.

## Declaration of Interests

The authors declare no competing interests.

## Author Contributions

Conceptualization, B.C.J, A.R., Z.A.K; Investigation, B.C.J, A.R., H.N, R.N., L.Q., T.L., J.Y.O., and O.K.B; Methodology, B.C.J, A.R., H.N., and Z.A.K.; Software, B.C.J and H.N.; Validation, B.C.J, A.R., H.N., Z.A.K.; Visualization B.C.J.; Project Administration and supervision, B.C.J and Z.A.K; Funding Acquisition, Z.A.K.; Writing (original, review, and editing), B.C.J and Z.A.K.

## Materials Availability

This study did not generate new unique reagents.

## Methods

### Animals

All procedures followed National Institutes of Health guidelines and were approved by the IACUC at the University of California, San Francisco. Mice were housed in a 12 h dark/light cycle (temperature and humidity controlled) with *ad libitum* water and PicoLab 5053 chow unless otherwise noted. Heterozygous *Calca*^Cre^ (Jax # 033168) or C57Bl/6J mice were used for all experiments. Experimental animals were over 8 weeks old and included both males and females, which were randomly assigned to control or experimental groups for behavioral studies. Mice with intragastric catheters or used in microendoscopic imaging studies were singly housed, and all other mice were group-housed when possible.

### Intracranial surgeries

*Calca*^Cre^ mice were anesthetized with 2% isofluorane and placed on a Kopf small animal stereotax. Viruses were pressure-injected into the parabrachial nucleus of *Calca*^Cre/+^ mice using glass capillaries and a Micro4 controller, at AP -5.1, ML ±1.5, DV -3.5 relative to bregma.

For retrograde tracing experiments, 0.15 µL pseudorabies-Introvert-GFP^17^ virus (undiluted, NIH CNNV from the Herpesvirus Core at Princeton University) was injected unilaterally and mice were perfused 2-4 days later.

For imaging experiments, mice were injected with 0.1-0.2 µL of an AAV expressing cre-dependent GCaMP6s (AAV1-CAG-FLEX-GCaMP6s-WPRE-SV40, Addgene 100842, 2.6-4.2E+12 gc/mL titer). During the same surgery, a commercially available GRIN lens from Inscopix (0.5 x 6.1 mm, 100-006647) was implanted at AP -5.0, ML 1.6, DV -3.6 or AP -5.2, ML 1.7, DV 3.7. Mice were baseplated 2-4 weeks after viral injection, and a subset of mice also received intragastric catheters as described below.

For behavioral experiments, mice were bilaterally injected with soma-tagged AA1-hSyn-DIO- GtACR2-FusionRed (Addgene 105677, 6.3E+12 gc/mL) for optogenetic experiments or AAV9- hSyn-DIO-hM4Di-mCherry (Custom virus from Boston Children’s Viral Core, 8.23E+12 gc/mL) for chemogenetic inhibition. Controls were transduced with AAV1-hSyn-DIO-mCherry (Addgene 50459, 5.7E+12 gc/mL). Custom-made optogenetic fibers were implanted at AP -5.2, ML ±1.6, DV 3.6 and all mice were given at least 2 weeks to recover before starting experiments. Animals with viral injections that missed on one or both sides of the brain were excluded.

### Intragastric catheterization surgery

Intragastric catheters were implanted as described^14,33^. Briefly, mice were anesthetized with isoflurane and sterile access catheters (C30PU-RGA1439, Instech Labs) were surgically implanted into the avascular stomach using a purse-string suture. The catheters were run under the skin to the back and attached to sterile vascular access buttons (VABM1B/22, Instech Labs). The buttons were secured under the skin above the scapulae of the mouse, with an externalized port was used to infuse solutions directly into stomach. An aluminum cap (VABM1C, Instech Labs) was placed over the port to protect it between experiments. Mice were given a minimum of one week to recover prior to the start of experiments.

### Histology

Mice were perfused with phosphate-buffered saline (PBS) and 30 mL of 10% formalin. Brains were post-fixed overnight then cryoprotected in 30% sucrose in PBS at 4°C, embedded in OCT, and stored frozen at -80C until processing. Brains were sectioned coronally at 30μm (every 3^rd^ section) on a cryostat. For determination of viral targeting, sections were imaged on a Zeiss LSM 510 confocal or Nikon Eclipse Ti2 widefield microscope.

For pseudorabies tracing, the pons, medulla, nodose and geniculate ganglia were collected. The ventral skull was transferred into PBS after overnight fixation and stored until ganglia dissection. All PRV-Introvert-GFP tissue underwent immunohistochemistry for GFP. Briefly, tissue was blocked in 2% normal donkey serum/PBST (0.1% Triton-X) and incubated in primary antibody (1:2000 chicken anti-GFP, Abcam ab13970). Brains were left in primary overnight, whereas ganglia were incubated for 72 h. Tissue was then washed 3 times in PBS and incubated in secondary antibody at room temperature for 1-3 h (1:000 donkey anti-chicken Alexa Fluor-488A, Sigma-Aldrich SAB4600031). Tissue was washed an additional 3 times in PBS, mounted, and stored at 4°C.

We performed whole-slide imaging of brain tissue using a Nikon Eclipse Ti2 widefield microscope and NIS Elements with a custom JOBS program. Sections were imaged at 10x (8- bit) with consistent exposure and gain, and images were exported as PNG files for QUINT. Ganglia were whole-mounted and imaged using a Zeiss LSM 510 confocal.

### QUINT Analysis

Pseudorabies tracing analysis was done using QUINT (https://quint-workflow.readthedocs.io/en/latest/QUINTintro.html)^57^. To register sections to an atlas, images were downsized to 10% using *Nutil*, and LUT settings were uniformly adjusted in ImageJ to optimize feature visualization. Sections were registered to the Allen Brain common coordinate framework using *QuickNII* and *Visualign*. For automated cell detection, images were downsized to 50% of their original size and pixel and object classifiers were created in *Ilastik* to identify GFP-positive cells. *Ilastik* output files underwent a Glasbey transformation using ImageJ, and these output files and the final *Visualign* files were quantified in *Nutil* using object splitting. Data is reported as the area of each nucleus analyzed that had GFP-positive signal (GFP pixel area/region area sampled = load). Regions included were manually verified by visual inspection to guarantee model accuracy, and the top 10 hits across brains were included. The NTS was manually split at the obex, with sections containing the obex included in the caudal category.

Regions with < 4000 pixels were filtered out, and the cerebellum and inferior and superior colliculi were excluded due to higher levels of background and damage. No cells were observed in these regions, although sparse labeling has been reported using monosynaptic rabies tracing^27^.

The *Calca*^Cre:GFP^ mouse line expresses nuclear-localized GFP that can be visualized with IHC. This endogenous GFP did not explain the expression pattern seen with PRV-Introvert-GFP, as we did not see GFP in regions of interest in *Calca*^Cre:GFP^ mice without virus injections (**Supplemental Fig. 2c**).

### Gastric emptying and distension

Fasted C57Bl/6J mice with IG catheters received a 1 mL infusion (saline, 20% intralipid, or Ensure) using a syringe pump (Harvard Apparatus, 70–2001) over 10 minutes (0.1 mL/min). Mice were perfused at the end of the infusion with 10 mL PBS and 10 mL formalin. During the perfusion, suture was used to ligate the pyloric sphincter. The GI tract was then dissected out, and additional sutures were placed 10 and 20 cm distal to the pyloric sphincter in the intestine. Infused solutions were not observed below 20 cm. GI segments were weighed before and after removal of contents. Content weight was normalized to body weight (g per 20 g body weight)^24^. Gastric emptying = (infused solution weight - stomach contents)/infused weight. Gross stomach area was measured from calibrated images in ImageJ.

### Pharmacology

All compounds were prepared in 0.9% sterile saline and stored at -20C prior to use, except for lithium chloride (LiCl), which was stored at 4°C. Compounds consisted of CCK octapeptide (10- 30 mg/kg; Tocris 1166), Exendin-4 (30-150 mg/kg; Tocris 1933), LiCl (84-100 mg/kg), deoxynivalenol (DON, 2.5 mg/kg; Tocris 3976), or DCZ (0.3mg/kg; Hellobio HB9126). All solutions were administered at a volume of 10µL/g BW.

### Ingestion Assays Optogenetics

Mice were habituated to bilateral fiber optic cables (Doric Lenses) before tests. A 473 nm laser delivered 0.8-1 mW of blue light per fiber either continuously or closed-loop (triggered by licks; laser off after 2 s without licking). Intake was monitored using lickometers in either a Coulborn Habitest Modular System or Med Associates Behavioral Chambers, which also controlled the laser. Sessions were excluded if the lickometer failed, mice did not engage (> 100 licks or < 10 min latency to lick), or the patch cord uncoupled.

For Ensure consumption experiments, mice either sated or fasted overnight were given 1-2h access to Ensure. For two-bottle choice tests, mice were water-deprived overnight and given access to water and either 0.5 mM quinine or 0.67 M glucose for 30 minutes, with and without continuous laser inhibition. Mice underwent two sessions for each experimental condition, and the order was counterbalanced across groups for the laser on and off condition. All solutions were familiar.

Meals were defined by an inter-meal interval (interval between licks) of > 5 minutes with a minimum intake of 5 licks (equivalent to ∼ 0.05 g Ensure). Bouts were separated by an inter-lick interval of 5 seconds and were at least 3 licks long. Data points for each animal are an average of 1-3 replicate sessions. For cumulative intake of Ensure, data was binned into 1 second intervals.

### Chemogenetics

For Ensure consumption experiments, mice were first habituated to Med Associates Behavioral Chambers with lickometers. Mice were given either saline or DCZ injections when placed in the behavior chambers, and then after 10 minutes were given 2 h access to Ensure. This was done for mice that were either sated or had been fasted overnight, and injections were counterbalanced.

For fast-refeed with solid food, mice were given either saline or DCZ and placed in a clean home cage. After 10 minutes, they were given a solid food pellet (AIN-76A, Research Diets), which was then hand-weighed at 30, 60, and 120 minute time points after access.

To measure daily baseline food intake, mice were placed in a cage where their food intake was tracked using FED3 devices (LabMaker)^58^. The FED3 devices were set to “free-feeding”, where Dustless Precision Pellets (Bio-Serv F0071) were monitored and a new pellet is dispensed 10 seconds after the previous pellet was removed. Mice were acclimated to using the FED3 as their food source for 3-5 days prior to the start of experiments, and saline or DCZ were administered 10-20 minutes before the start of the dark cycle. Sessions were excluded if the FED device jammed or if mice did not start eating during the first 30 minutes of the dark cycle.

For meal analysis of FED3 data, pellet number and the inter-pellet-intervals were used. Bouts were assumed to be each pellet and we were unable to look at this as an independent metric. Meal definitions were the same as for Ensure, with meal defined by an inter-pellet interval of > 5 minutes with a minimum intake of 3 pellets (0.06 g food).

### Conditioned taste avoidance

For optogenetic experiments, water restricted mice underwent 4 training days with a 30-minute one-bottle session each day, followed by 3 days of 30-minute two-bottle choice tests. Training sessions alternated between access to water and sucralose (0.8 mg/ml sucralose in saline).

Sucralose was paired with continuous laser inhibition followed by 100 mg/kg IP LiCl with no laser inhibition, whereas water days had no laser or injection. Two bottle- choice tests were between water and sucralose, and laser inhibition only occurred on day 2 of the two-bottle tests. All mice also received daily 2 h water access in their home cage to provide additional hydration throughout the experiment.

For chemogenetic experiments, mice received 0.3 mg/kg injections of DCZ 10 minutes prior to sucralose access on training days, and only one two-bottle choice assay was performed to measure sucralose avoidance.

For CTA to IG infusions, C57Bl/6j mice were used. Sucralose access was paired with subsequent IG infusions (1 mL intralipid, Ensure, or saline, and 1.5 mL Ensure at a rate of 0.1 mL/min) instead of LiCl injections. Some mice went through two rounds of training with different IG infusion pairings, where in the second test the sucralose was flavored with orange kool-aid.

### Microendoscopic imaging

#### Data Acquisition

Microendoscopic videos were acquired using Inscopix acquisition software (v151) at 20Hz, 8 gain, 0.4-0.8 mm^−2^ 455 nm LED power, and 2x spatial downsampling. Data was recorded using nVista (v.3.0), nVoke (v2.0) or nVue (1.0, 2.0) systems. Prior to recording, mice were connected to cameras with the LED on for 10 minutes for habituation and stabilization of GCaMP6s signal. A baseline of 10 minutes was collected prior to all manipulations. In some cases, mice were head-fixed to stabilize the recordings.

For self-paced licking experiments, mice were fasted overnight and placed in Coulborn chambers, and licking for various solutions (saline, 0.24 g/mL glucose, 20% Intralipid, 0.8 mg/mL sucralose in saline, and Ensure) was tracked using a custom made capacitive lickometer connected to the Inscopix acquisition system.

Intermittent Ensure access experiments were performed in a Davis Rig (MED-DAV-160M, Med Associates), where mice underwent three days of training and habituation prior to experiments^13^. For recordings, fasted mice were given repeating 60 s access to Ensure, with 120 s breaks between access trials, for a total of 15 trials over the course of 60 minutes. For IG infusions, mice were fasted overnight and received 1 or 1.5 mL delivery of Ensure, 20% Intralipid, 24% glucose, 45% Cherry-flavored Proteinex, or saline. Hormones were administered intraperitonially (IP) in fasted mice.

For solid food experiments, mice were fasted overnight and given 55-60 min access to their normal chow (5053 PicoLab) or high-fat diet (60%, Research Diets, D12492). Some of these recordings occurred in the homecage, and some in Coulborn chambers. Mice were recorded throughout the recording, and bites of food were hand-scored using BORIS software. Mice all had previous exposure to the high-fat diet prior to experiments.

For experiments comparing responses to Ensure and aversive agents, mice were given Ensure access for 10 minutes and then received either 84 mg/kg IP LiCl or 2.5 mg/kg DON after a 10- minute break. LiCl was paired with vanilla Ensure, and DON was paired with chocolate Ensure.

#### Video processing

Videos were processed using Inscopix Data-Processing Software (v1.9.4). Videos were spatially (factor 5) and temporally (factor 2) binned to 4 Hz, spatially band-pass filtered, and motion corrected. To further reduce residual motion, videos were subjected to an additional round of motion correction consisting of rigid registration followed by piecewise non-rigid registration^59^, with parameters empirically optimized for each dataset. Videos with uncorrectable motion were discarded. In some cases, neurons within the videos had two distinct motion patterns, and two motion-corrected versions of the videos were created using ROIs around specific subsets of neurons. Large motion artifacts were excised. Activity traces for individual neurons were extracted using the constrained non-negative matrix factorization (CNMF-E) pipeline built into the Inscopix software, with output units of change in fluorescence over noise. Neurons were manually refined to remove duplicates, motion artifacts, and over-segmentation. Neural traces from 4-10 mice per experiment were pooled for analysis.

### Data Analysis

#### Self-paced feeding

For self-paced feeding experiments, neural traces were z-scored to the 10-minute baseline prior to food access. Mean activity was then averaged over the first 5 and the first 10 minutes of solution access to capture differences in temporal distribution of intake over time. Neurons with mean *z* > +1 in either epoch were classified as activated, and those with *z* < -1 were classified as inhibited. Neural responses across the first 10 minutes were used to compare responses between solutions.

For extended solid food recordings, the z-score of the last 10 minutes (45-55 or 50-60 minutes) was calculated and compared to responses at the start of food access. Neurons meeting specific criteria were further classified into response types: Neurons identified as described above had “rapid” responses to food ingestion. “Delayed” neurons exhibited z > +1 (or < -1 for inhibited neurons) in the last 10 minutes but not the first 10 minutes. By definition, all delayed neurons also show ramping (a difference of > +1 between the first and last 10 minutes), though there were a small number neurons also identified as ramping in the “Rapid” category (<2%).

Activation or inhibition of neurons across the duration of the recording was quantified by counting neurons with mean z-score > +1 (or < -1) in each 5-minute epoch.

#### Intermittent Access

For intermittent access experiments, activity was averaged over each 60-s access period. Neurons were categorized as activated if mean z ≥ +1 in more than 50% of trials. The consistency of activated neural responses were the percentage of trials where the mean response was ≥ 1 *z*. “Start” and “end” values are the average z during the first and last three trials containing ≥ 3 licks, normalized to lick count.

#### Bout-Based Analysis

Bout activity was calculated as the mean z-score across all bouts during the sessions (30-60 min), normalized to lick count where indicated. Interbout activity was calculated from the same traces after masking out bout periods. In some analyses, interbout activity was further split into the first and last 10 min of the session. Bouts were defined in the same way as in behavioral assays.

To analyze single-cell responses to licking or bites, neuronal traces were z-scored to the 10-s baseline immediately preceding the first lick in each bout, and responses were quantified over the subsequent 0-20 s. For bout-offset analysis, traces were re-z-scored to the 10 s preceding the final lick in a bout and quantified over the next 0-20 second. Only bouts ≥ 10 s in duration and separated by ≥ 10 s were analyzed to guarantee independent baselines. All qualifying bouts from a given neuron were averaged to yield one PSTH per cell. PSTHs were smoothed with a moving average window equal to one-third of the 0-20 s response window, and time to 50% of the peak (rise) or trough (decay) was determined.

To quantify the relationship between consummatory behavior (bouts) and neural activity, z-scored traces (Gaussian-smoothed, σ = 2 s) were regressed onto a binary bout-state regressor (1 during a lick bout, 0 during inter-bout intervals) using ordinary least squares, and the coefficient of determination (R²) was used to quantify variance explained. To confirm that observed R² values reflected genuine neural-behavioral coupling, a circular-shift null distribution was generated by shifting the bout-state vector by 500 independently drawn random offsets (minimum 30 s) and computing R² for each shift. Observed R² was compared against the mean shuffle R² using a paired one-tailed t-test across neurons.

#### IG Infusions

Following IG infusions, neurons were classified as activated if they passed two criteria. First, that mean z ≥ +1 during the 10-15 min infusion or in the 10 min after the infusion. Second, that z-score was ≥ 1 more than 70% of the time during a sliding 60 second window. This captures sustained but not necessarily continuous responses and excludes brief transients. The first frame of that qualifying window was defined as the response start time for each cell. Survival curves were generated using response times to show the fraction of activated neurons that came online over time.

#### Aversive Agents

For Ensure vs. aversive agents, Ensure responses were z-scored to the 10-min pre-access baseline. LiCl and DON responses were z-scored to the 8-min pre-injection baseline and scored with the same 60-s sliding window rule used for IG infusions.

#### Graphing

Heatmaps were downsampled to 1 hz, and neural traces were smoothed by a moving average filter for 20 data points for graphing purposes only. Raw data were used for all quantitative analyses.

## Statistical Analysis

Data are shown as mean ± s.e.m. (error bars or shaded areas). For behavioral tests, sample size is the number of mice per group. For imaging, sample size is the number of neurons.

Graphing and analyses were performed in GraphPad Prism 10 (GraphPad Software) or using custom scripts in MATLAB R2025b.

A summary of statistical tests used in all figures is summarized in **Supplemental Table 1**. Normality was assessed with Shapiro-Wilk. For two-group comparisons, two-tailed Student’s t-tests were conducted with parametric data and Mann Whitney *U* tests with non-parametric data (paired or unpaired where appropriate). Air- versus Ensure-bout distributions (**Fig. 1g**) and mean IG and post-IG Ensure response distributions (**Fig. 3h**) were compared with Kolmogorov– Smirnov tests. Lick count versus response magnitude was evaluated by simple linear regression. Log-rank (Mantel-Cox) tests were used to compare survival curves showing neuron response times. Comparisons across multiple groups were analyzed using one-way ANOVA (Holm-Šidák’s post hoc) or Kruskal Wallis (Dunn post hoc). Šidák’s multiple comparisons were used with two-way ANOVA.

Pairwise odds ratios for cell activation were obtained from 2×2 contingency tables for each condition pair (activated vs. not activated) and applying Fisher’s exact test to estimate odds ratios with 95% confidence intervals. *P*-values were adjusted for multiple comparisons using a Bonferroni correction. Power calculations were used to predetermine sample size when prediction of effect size was possible. Experiments were not randomized and investigators were not formally blinded.

---

**Supplemental Figure 1.**
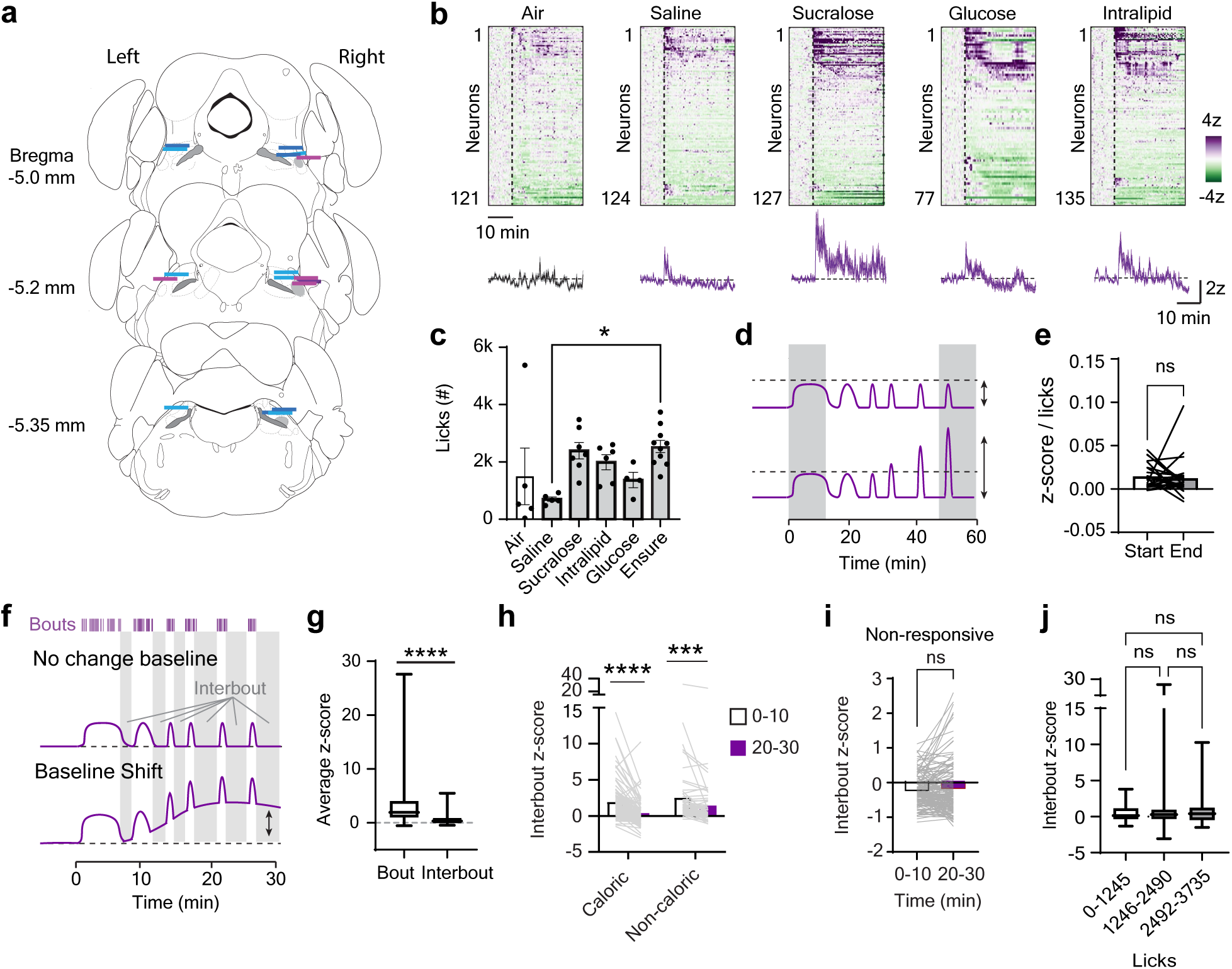
Analysis of CGRP neuron responses to ingestion. **a**, Approximate GRIN lens placements for all animals used for single-cell imaging. Blue lines are in the more dorsal orientation, and purple lines were categorized as lateral. **b**, Heatmaps and averaged response of neurons to self-paced licking for various solutions. **c**, The amount mice licked for different solutions or air during single-cell imaging experiments. **d**, Model for how CGRP neurons responses to ingestion could change over time as food consumption progresses. **e**, Mean z-scored responses normalized to number of licks for the first (start) and last (end) three trials of the intermittent access test. **f**, Model for looking at changes in tonic CGRP neuron activity (baseline shift) during food consumption. **g**, There were greater CGRP neural responses during licking bouts compared to interbout intervals for Ensure. **h**, Mean interbout z-score of activated neurons at the start and end of a licking session (first vs. last 10 minutes). **i**, Average interbout z-scored response in the first and last 10 minutes of Ensure access of neurons categorized as “non-responsive”. **j**, Interbout z-score of activated neurons in the last 10 minutes of a licking session for caloric solutions (Ensure, glucose, and Intralipid), binned based on the total licks consumed by the end of the session. ns mean not significant. **** *p* < .0001, *** *p* < .001, ** *p* < .01. * *p* < .05

**Supplemental Figure 2.**
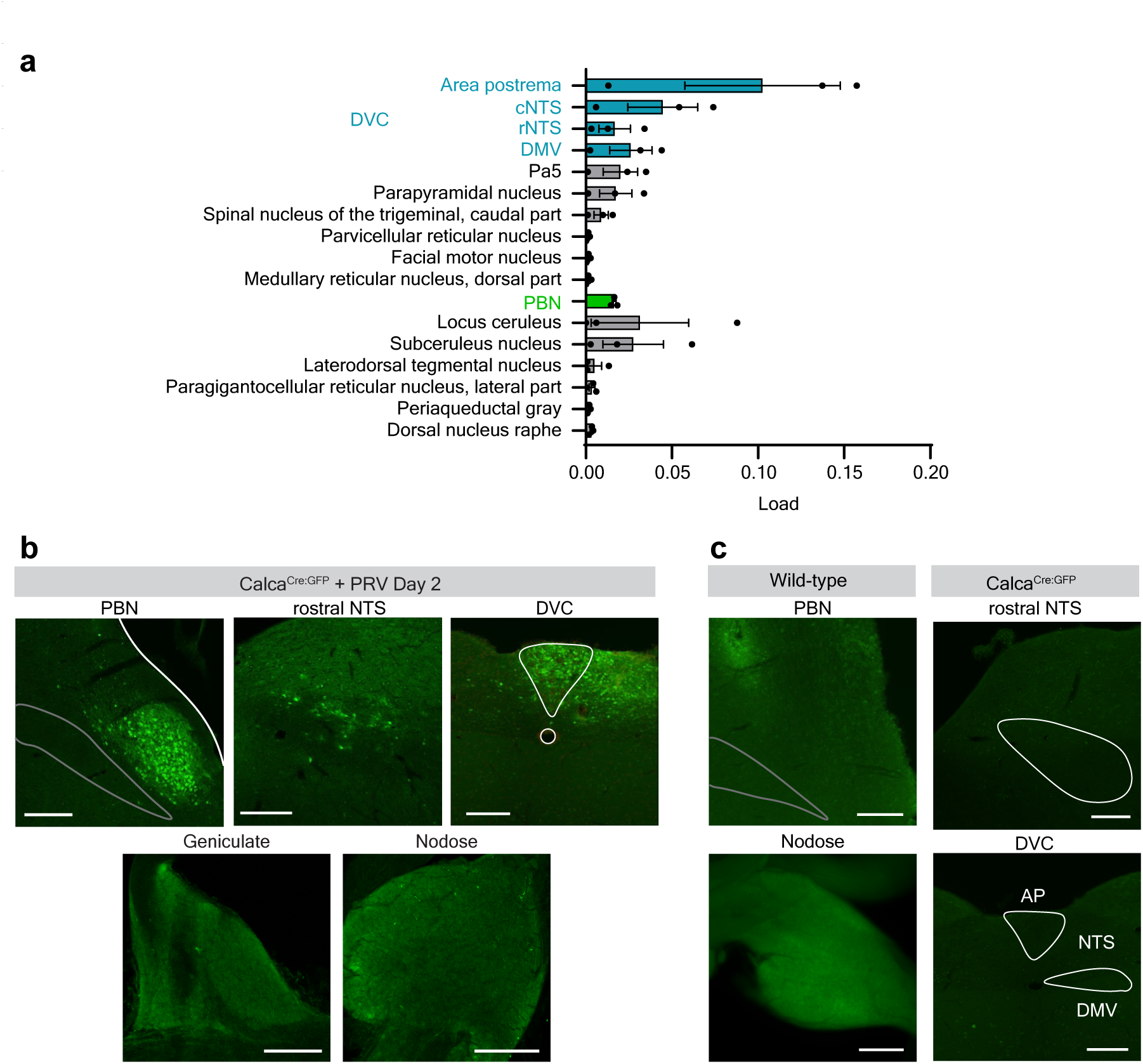
Characterization of PRV-Introvert-GFP and brainstem inputs to CGRP neurons. **a**, Quantification expression of PRV-Introvert-GFP on day 3 following injection into CGRP neurons in the parabrachial nucleus (*N* = 3 mice). GFP expression was calculated using QUINT, and is reported as GFP-positive pixels / total area sampled per nucleus (load). **b**, Example images of PRV-Introvert-GFP on day 2 following injection, showing expression in the PBN and dorsal vagal complex (DVC), but not the geniculate or nodose ganglia. **c**, Controls showing that injection of PRV-Introvert-GFP into wild-type mice does not lead to GFP expression in either the PBN or nodose ganglia, and endogenous expression of GFP in the Calca^Cre:GFP^ mouse line is not present in the NTS and surrounding regions. scale bars, 200 um. AP, area postrema. NTS, nucleus of the solitary tract. DMV, dorsal motor vagus.

**Supplemental Figure 3.**
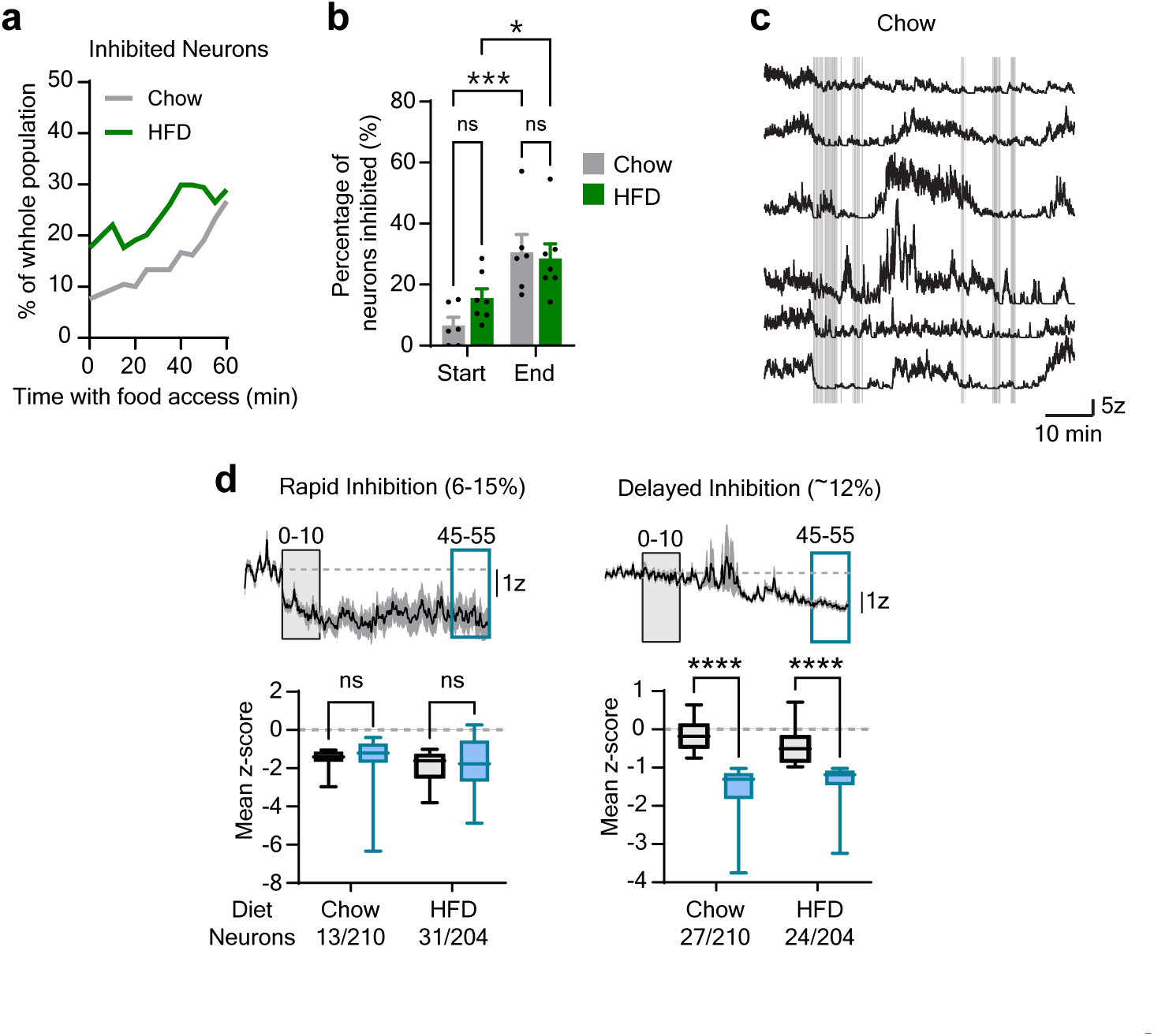
A subset of CGRP neurons are inhibited by ingestion. **a**, Percentage of neurons that were inhibited in 5-min bins over the course of 55 min of food access. **b**, Quantification of the number of inhibited neurons in each mouse that was recorded from during the first and last 5 minutes of the recording session. **c**, Individual traces of neurons that are inhibited by intake of chow. Vertical gray bars are bites. **d**, Top, average neural traces and bottom, mean z response during the first and last 10 minutes epochs of food access of neurons that were inhibited when first given food access (rapid) or at the end of the session (delayed). **** *p* < .0001, ** *p* < .001, * *p* < .05.

**Supplemental Figure 4.**
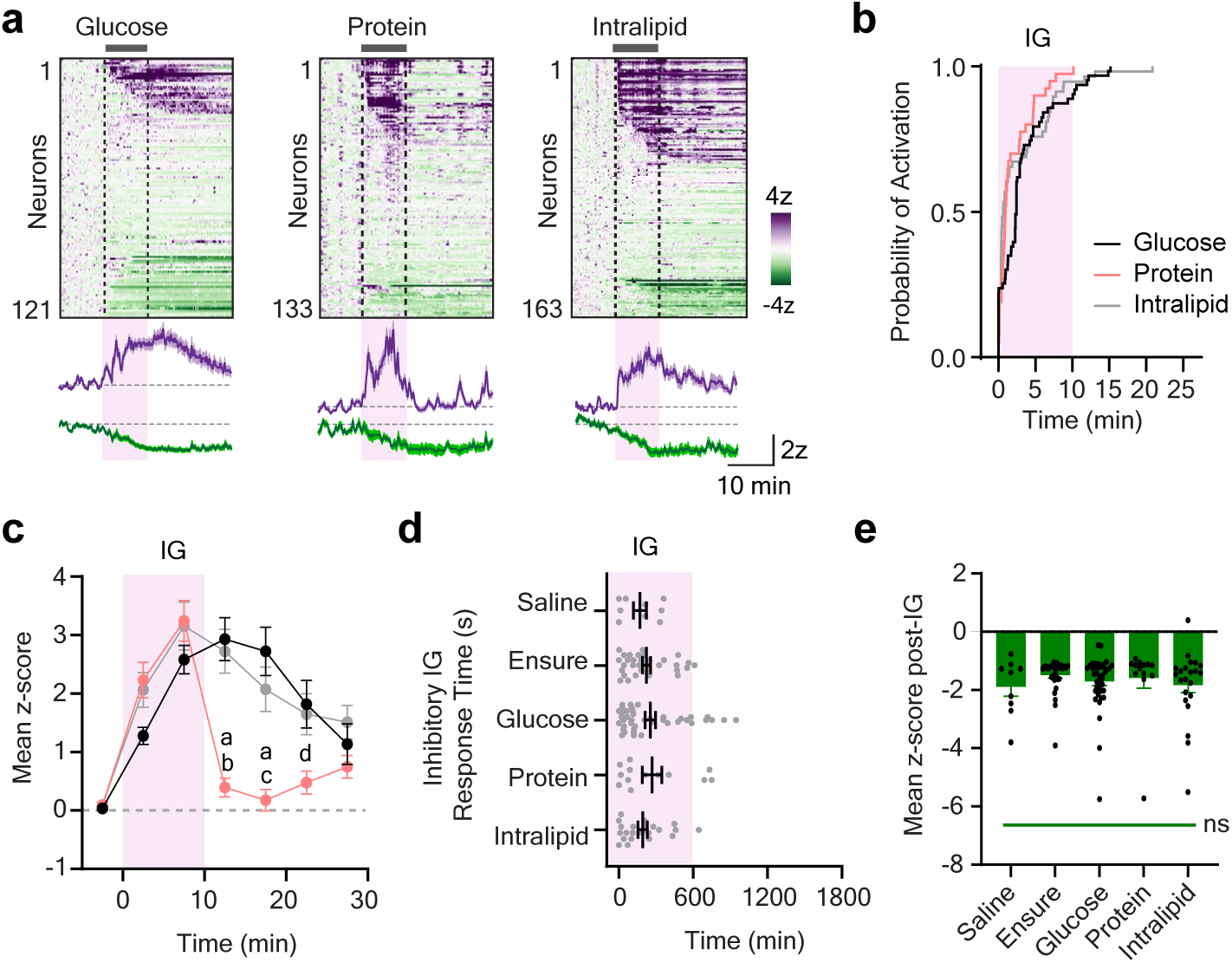
Concentrated IG macronutrients alter CGRP neuron activity. **a**, Heatmaps of CGRP neural responses to intragastic (IG) infusions of 1 mL of 24% glucose, 16% protein, or 20% Intralipid. **b**, Cumulative activation of CGRP neurons over time responding to IG infusions of each macronutrient. Each step represents neurons reaching activation threshold. Log-rank test, p = 0.17. **c,** Mean z-score of activated neurons during 5 min epochs across the duration of the experiment in response to macronutrient infusions. **d**, Start of the inhibitory response for each neuron inhibited by the IG infusions. **e**, Mean inhibitory response during the 10 minutes post-IG. ns, not significant. a, *p* <.0001 compared to glucose b, *p* < .0001 compared to Intralipid c, *p* <.001 compared to Intralipid d, *p* < .05 compared to glucose and Intralipid.

**Supplemental Figure 5.**
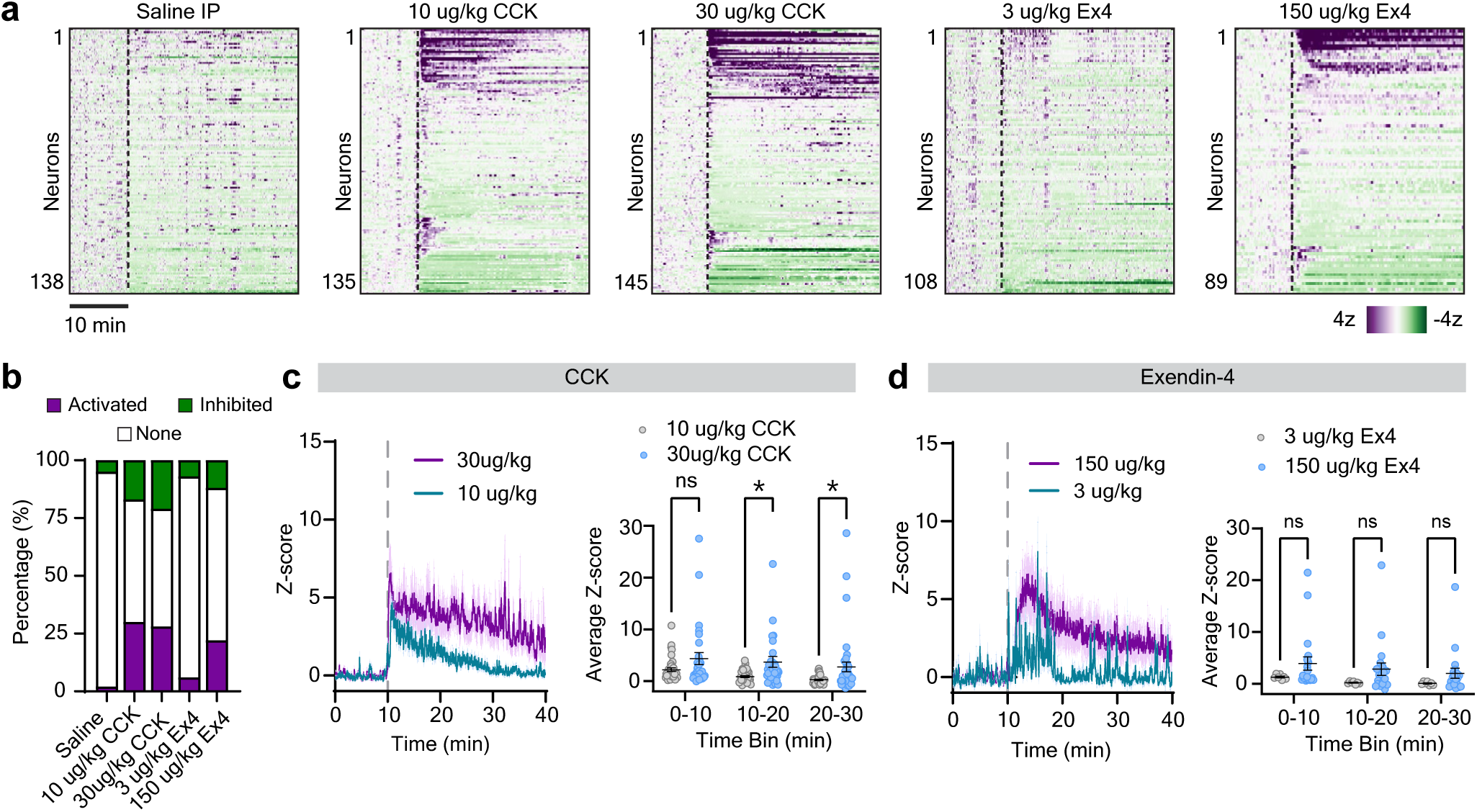
Gut satiety hormones activate CGRP neurons. **a**, Heatmaps of CGRP neural responses to i.p. injections of saline or different doses of CCK and the GLP-1 agonist, Exendin-4 (Ex4). **b**, Quantification of the neural responses following injections. **c**, Left, average response of the activated neurons to a low and high does of CCK. Right, quanitification of the mean z-scored response during different time bins following CCK injections showed that neurons had more sustained responses to 30ug/kg CCK, specifically during the 10-20 and 20-30 minute time bins. **d**, Left, Averaged trace showing response of activated neurons to two doses of Exendin-4. Right, there were no significant differences in responses over time between the Ex4 doses. ns mean not significant. * *p* < .05.

**Supplemental Figure 6.**
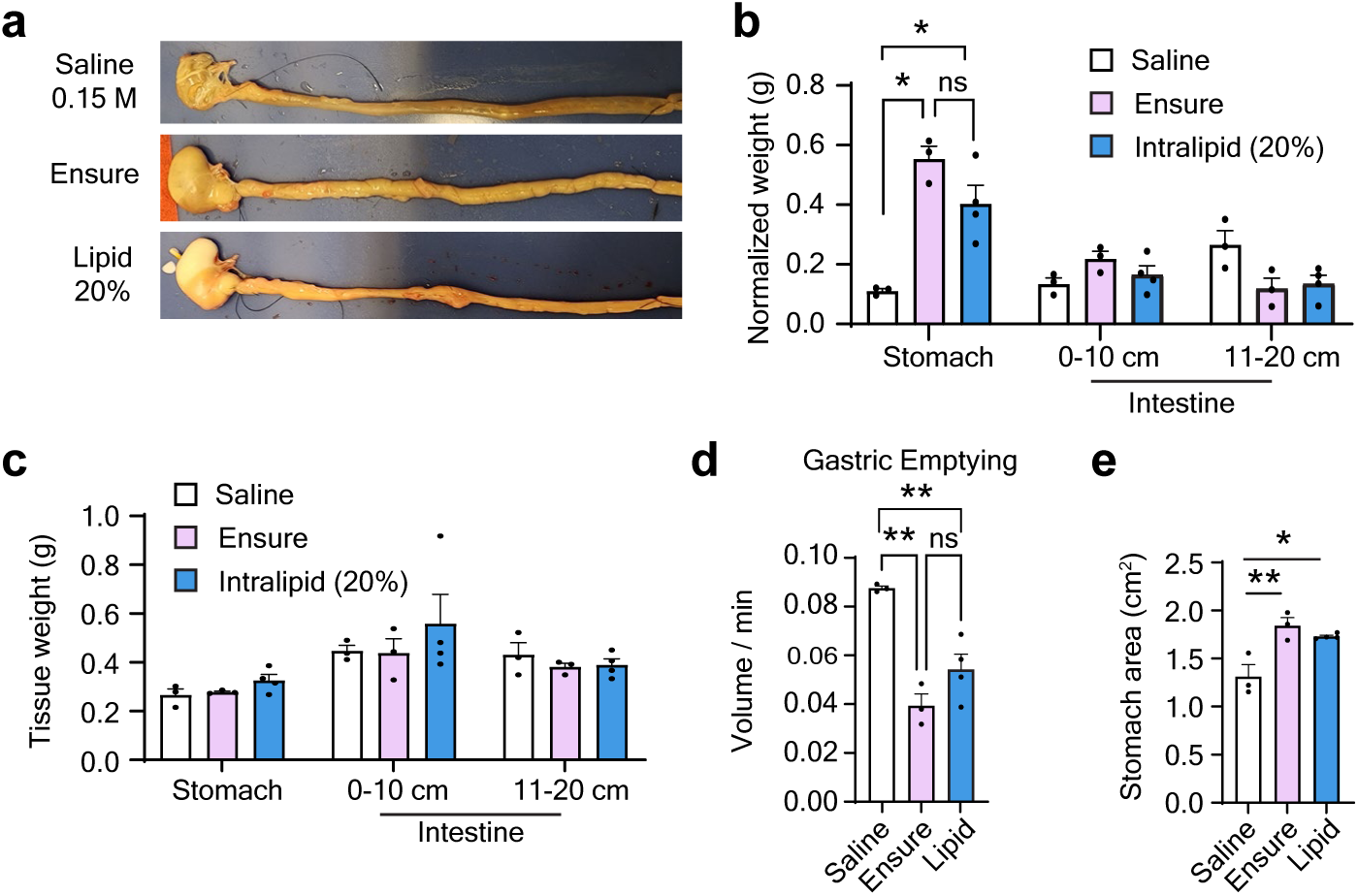
Gastrointestinal changes following IG infusions. **a**, Mice received a 1 mL intragastric infusions of saline, Ensure, or intralipid. At the end of the infusion, gastrointestinal tissue was collected and contents were analyzed. **b**, Weight of the contents of the stomach and different sections of the intestines. **c**, No differences in the weight of GI tissue (without contents) were observed between mice. **d**, Gastric emptying was significantly slower following infusion of Ensure or intralipid than saline. **e**, Surface area of the stomach was greater following infusion of Ensure and intralipid compared to saline. ns means not significant. * *p* < .05, ** *p* <.01.

**Supplemental Figure 7.**
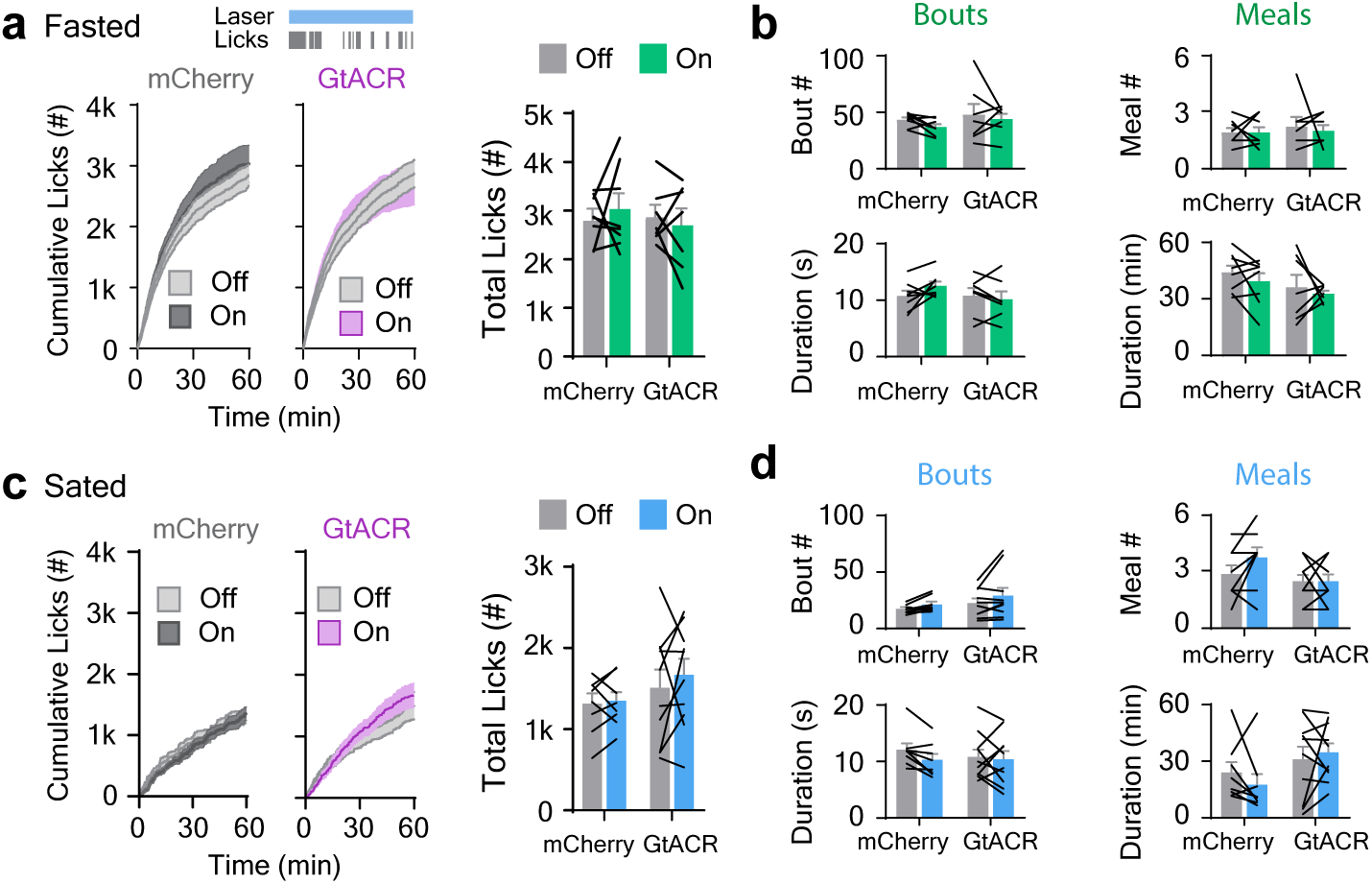
Tonic optogenetic inhibition of CGRP neurons has no effect on liquid food intake. Continuous bilateral inhibition of CGRP-PBN neurons during self-paced intake of Ensure. There was no effect on mice expressing either mCherry or GtACR on **a**, cumulative rate or total licks or **b**, bout and meal number and duration in mice that had been fasted overnight. There was also no effect on **c**, total licks or **d**, bout and meal number and duration in sated mice.

**Supplemental Figure 8.**
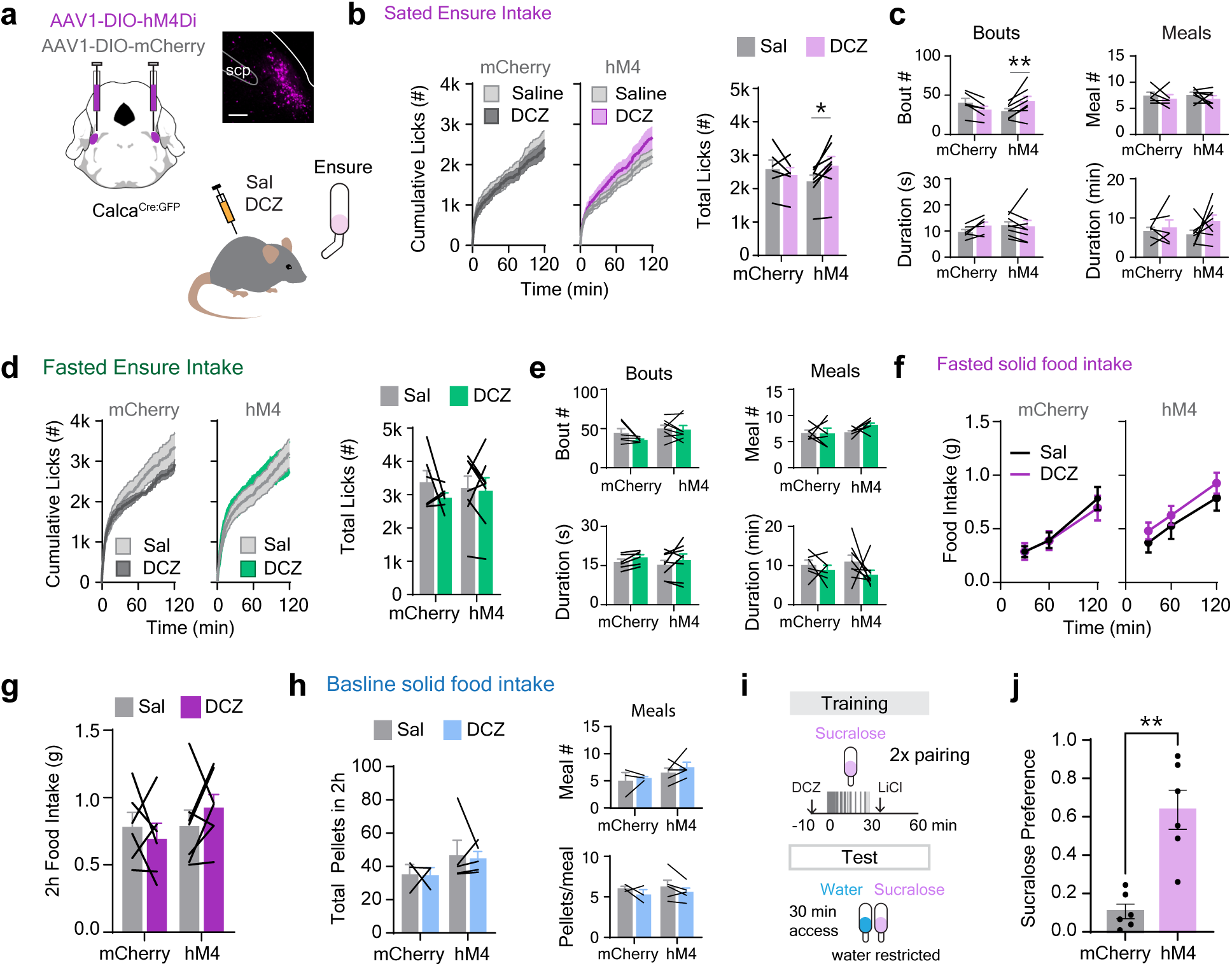
Effects of chemogenetically silencing CGRP neurons. **a**, Left, *Calca*^Cre:GFP^ mice were injected with AAV1-DIO-hM4Di or AAV1-DIO-mCherry for tonic silencing of CGRP-PBN neurons. Right, image of hM4Di expression and schematic of IP saline (Sal) or DCZ injection while monitoring Ensure intake. **b**, Silencing CGRP neurons increased Ensure intake in fed mice by **c**, increasing the number of bouts but not bout duration, meal number, or meal duration. **d**, DCZ did not alter cumulative intake, total licks, or **e**, bout or meal microstructure in mice fasted overnight. **f**, There was no effect of silencing CGRP neurons on cumulative or **g** total intake of solid food after an overnight fast. **h**, FED3 devices were used to monitor the first 2h of daily food intake, and DCZ had no effect on total intake, meal number or meal size. **i**, Training and testing paradigm for conditioned taste avoidance, where CGRP neurons were inhibited during training using DCZ. **j**, Mice expressing hM4D developed an attenuated CTA compared to control mice. scale bar 200 um, scp, superior cerebellar peduncle. \**p* < .05, \*\**p* < .01

## Notes

### Competing Interest Statement

The authors have declared no competing interest.

### Summary of Updates

Supplemental Figure 8 and Table 1 added. Text revised to discuss new data.

## References

1. Carter, M.E., Soden, M.E., Zweifel, L.S., and Palmiter, R.D. (2013). Genetic identification of a neural circuit that suppresses appetite. Nature 503, 111–114. 10.1038/nature12596.

2. Campos, C.A., Bowen, A.J., Han, S., Wisse, B.E., Palmiter, R.D., and Schwartz, M.W. (2017). Cancer-induced anorexia and malaise are mediated by CGRP neurons in the parabrachial nucleus. Nat. Neurosci. 20, 934–942. 10.1038/nn.4574.

3. Campos, C.A., Bowen, A.J., Schwartz, M.W., and Palmiter, R.D. (2016). Parabrachial CGRP Neurons Control Meal Termination. Cell Metab. 23, 811–820. 10.1016/J.CMET.2016.04.006.

4. Chen, J.Y., Campos, C.A., Jarvie, B.C., and Palmiter, R.D. (2018). Parabrachial CGRP Neurons Establish and Sustain Aversive Taste Memories. Neuron 100, 891–899.e5. 10.1016/j.neuron.2018.09.032.

5. Han, S., Soleiman, M.T., Soden, M.E., Zweifel, L.S., and Palmiter, R.D. (2015). Elucidating an Affective Pain Circuit that Creates a Threat Memory. Cell 162, 363–374. 10.1016/j.cell.2015.05.057.

6. Carter, M.E., Han, S., and Palmiter, R.D. (2015). Parabrachial Calcitonin Gene-Related Peptide Neurons Mediate Conditioned Taste Aversion. J. Neurosci. 35, 4582–4586. 10.1523/JNEUROSCI.3729-14.2015.

7. Pyeon, G.H., Cho, H., Chung, B.M., Choi, J.-S., and Jo, Y.S. (2025). Parabrachial CGRP neurons modulate active defensive behavior under a naturalistic threat. eLife 14, e101523. 10.7554/eLife.101523.

8. Smith, J.A., Ji, Y., Lorsung, R., Breault, M.S., Koenig, J., Cramer, N., Masri, R., and Keller, A. (2023). Parabrachial Nucleus Activity in Nociception and Pain in Awake Mice. J. Neurosci. 43, 5656–5667. 10.1523/JNEUROSCI.0587-23.2023.

9. Kang, S.J., Liu, S., Ye, M., Kim, D. Il, Pao, G.M., Copits, B.A., Roberts, B. Z., Lee, K.F., Bruchas, M.R., and Han, S. (2022). A central alarm system that gates multi-sensory innate threat cues to the amygdala. Cell Rep. 40, 111222. 10.1016/j.celrep.2022.111222.

10. Zimmerman, C.A., Bolkan, S.S., Pan-Vazquez, A., Wu, B., Keppler, E.F., Meares-Garcia, J.B., Guthman, E.M., Fetcho, R.N., McMannon, B., Lee, J., et al. (2025). A neural mechanism for learning from delayed postingestive feedback. Nature, 1–10. 10.1038/s41586-025-08828-z.

11. Palmiter, R.D. (2018). The Parabrachial Nucleus: CGRP Neurons Function as a General Alarm. Trends Neurosci. 41, 280–293. 10.1016/j.tins.2018.03.007.

12. Campos, C.A., Bowen, A.J., Roman, C.W., and Palmiter, R.D. (2018). Encoding of danger by parabrachial CGRP neurons. Nature 555, 617–622. 10.1038/nature25511.

13. Aitken, T.J., Liu, Z., Ly, T., Shehata, S., Sivakumar, N., Medina, N.L.S., Gray, L.A., Zhang, J., Dundar, N., Barnes, C., et al. (2024). Negative feedback control of hypothalamic feeding circuits by the taste of food. Neuron 112, 3354–3370.e5. 10.1016/j.neuron.2024.07.017.

14. Ly, T., Oh, J.Y., Sivakumar, N., Shehata, S., La Santa Medina, N., Huang, H., Liu, Z., Fang, W., Barnes, C., Dundar, N., et al. (2023). Sequential appetite suppression by oral and visceral feedback to the brainstem. Nature C24, 130–137. 10.1038/s41586-023-06758-2.

15. Wang, H., Lou, R., Wang, Y., Hao, L., Wang, Q., Li, R., Su, J., Liu, S., Zhou, X., Gao, X., et al. (2025). Parallel gut-to-brain pathways orchestrate feeding behaviors. Nat. Neurosci. 28, 320–335. 10.1038/s41593-024-01828-8.

16. Kim, D.-Y., Heo, G., Kim, M., Kim, H., Jin, J.A., Kim, H.-K., Jung, S., An, M., Ahn, B.H., Park, J.H., et al. (2020). A neural circuit mechanism for mechanosensory feedback control of ingestion. Nature 580, 376–380. 10.1038/s41586-020-2167-2.

17. Pomeranz, L.E., Ekstrand, M.I., Latcha, K.N., Smith, G.A., Enquist, L.W., and Friedman, J.M. (2017). Gene Expression Profiling with Cre-Conditional Pseudorabies Virus Reveals a Subset of Midbrain Neurons That Participate in Reward Circuitry. J. Neurosci. 37, 4128–4144. 10.1523/JNEUROSCI.3193-16.2017.

18. Borison, H.L. (1989). Area Postrema: Chemoreceptor circumventricular organ of the medulla oblongata. Prog. Neurobiol. 32, 351–390. 10.1016/0301-0082(89)90028-2.

19. Miller, A.D., and Leslie, R.A. (1994). The Area Postrema and Vomiting. Front. Neuroendocrinol. 15, 301–320. 10.1006/frne.1994.1012.

20. Zhang, C., Kaye, J.A., Cai, Z., Wang, Y., Prescott, S.L., and Liberles, S.D. (2021). Area Postrema Cell Types that Mediate Nausea-Associated Behaviors. Neuron 109, 461–472.e5. 10.1016/j.neuron.2020.11.010.

21. Blomquist, A.J., and Antem, A. (1965). Localization of the terminals of the tongue afferents in the nucleus of the solitary tract. J. Comp. Neurol. 124, 127–130. 10.1002/cne.901240110.

22. May, O.L., and Hill, D.L. (2006). Gustatory terminal field organization and developmental plasticity in the nucleus of the solitary tract revealed through triple-fluorescence labeling. J. Comp. Neurol. 497, 658–669. 10.1002/cne.21023.

23. Norgren, R., and Leonard, C.M. (1973). Ascending central gustatory pathways. J. Comp. Neurol. 150, 217–237. 10.1002/cne.901500208.

24. Bai, L., Mesgarzadeh, S., Ramesh, K.S., Huey, E.L., Liu, Y., Gray, L.A., Aitken, T.J., Chen, Y., Beutler, L.R., Ahn, J.S., et al. (2019). Genetic Identification of Vagal Sensory Neurons That Control Feeding. Cell 179, 1129–1143.e23. 10.1016/j.cell.2019.10.031.

25. Lowenstein, E.D., Ruffault, P.-L., Misios, A., Osman, K.L., Li, H., Greenberg, R.S., Thompson, R., Song, K., Dietrich, S., Li, X., et al. (2023). Prox2 and Runx3 vagal sensory neurons regulate esophageal motility. Neuron 111, 2184–2200.e7. 10.1016/j.neuron.2023.04.025.

26. Roman, C.W., Derkach, V.A., and Palmiter, R.D. (2016). Genetically and functionally defined NTS to PBN brain circuits mediating anorexia. Nat. Commun. 7, 11905. 10.1038/ncomms11905.

27. Korkutata, M., De Luca, R., Fitzgerald, B., Khanday, M.A., Arrigoni, E., and Scammell, T.E. (2025). Afferent Projections to the Calca/CGRP-Expressing Parabrachial Neurons in Mice. J. Comp. Neurol. 533, e70018. 10.1002/cne.70018.

28. Kim, J.H., Kromm, G.H., Barnhill, O.K., Sperber, J., Heuer, L.B., Loomis, S., Newman, M.C., Han, K., Gulamali, F.F., Legan, T.B., et al. (2022). A discrete parasubthalamic nucleus subpopulation plays a critical role in appetite suppression. eLife 11, e75470. 10.7554/eLife.75470.

29. Sabatini, P.V., Frikke-Schmidt, H., Arthurs, J., Gordian, D., Patel, A., Rupp, A.C., Adams, J.M., Wang, J., Beck Jørgensen, S., Olson, D.P., et al. (2021). GFRAL-expressing neurons suppress food intake via aversive pathways. Proc. Natl. Acad. Sci. 118, e2021357118. 10.1073/pnas.2021357118.

30. Simon, S.A., de Araujo, I.E., Gutierrez, R., and Nicolelis, M.A.L. (2006). The neural mechanisms of gustation: a distributed processing code. Nat. Rev. Neurosci. 7, 890–901. 10.1038/nrn2006.

31. Berthoud, H.-R., and Neuhuber, W.L. (2000). Functional and chemical anatomy of the afferent vagal system. Auton. Neurosci. 85, 1–17. 10.1016/S1566-0702(00)00215-0.

32. Terrier, L.-M., Hadjikhani, N., and Destrieux, C. (2022). The trigeminal pathways. J. Neurol. 269, 3443–3460. 10.1007/s00415-022-11002-4.

33. Beutler, L.R., Chen, Y., Ahn, J.S., Lin, Y.-C., Essner, R.A., and Knight, Z.A. (2017). Dynamics of Gut-Brain Communication Underlying Hunger. Neuron 96, 461–475.e5. 10.1016/j.neuron.2017.09.043.

34. Ueno, A., Lazaro, R., Wang, P.-Y., Higashiyama, R., Machida, K., and Tsukamoto, H. (2012). Mouse intragastric infusion (iG) model. Nat. Protoc. 7, 771–781. 10.1038/nprot.2012.014.

35. Sternson, S.M., and Eiselt, A.-K. (2017). Three Pillars for the Neural Control of Appetite. Annu. Rev. Physiol. 79, 401–423. 10.1146/annurev-physiol-021115-104948.

36. Talsania, T., Anini, Y., Siu, S., Drucker, D.J., and Brubaker, P.L. (2005). Peripheral Exendin-4 and Peptide YY3–36 Synergistically Reduce Food Intake through Different Mechanisms in Mice. Endocrinology 146, 3748–3756. 10.1210/en.2005-0473.

37. Mack, C.M., Moore, C.X., Jodka, C.M., Bhavsar, S., Wilson, J.K., Hoyt, J.A., Roan, J.L., Vu, C., Laugero, K.D., Parkes, D.G., et al. (2006). Antiobesity action of peripheral exenatide (exendin-4) in rodents: effects on food intake, body weight, metabolic status and side-effect measures. Int. J. Obes. 30, 1332–1340. 10.1038/sj.ijo.0803284.

38. Ding, H., Wang, M., Zhang, J., Wan, C., Huang, Z., Liu, L., and Liu, J. (2025). Exendin-4 induced retching-like behavior mediated by postsynaptic effect via AMPA receptors in the area postrema of mice. Am. J. Physiol.-Endocrinol. Metab. 329, E254–E265. 10.1152/ajpendo.00174.2025.

39. Welzl, H., D’Adamo, P., and Lipp, H.-P. (2001). Conditioned taste aversion as a learning and memory paradigm. Behav. Brain Res. 125, 205–213. 10.1016/S0166-4328(01)00302-3.

40. Patel, A.R., Frikke-Schmidt, H., Sabatini, P.V., Rupp, A.C., Sandoval, D.A., Myers, M.G., and Seeley, R.J. (2023). Neither GLP-1 receptors nor GFRAL neurons are required for aversive or anorectic response to DON (vomitoxin). Am. J. Physiol. Regul. Integr. Comp. Physiol. 10.1152/ajpregu.00189.2022.

41. Jarvie, B.C., Chen, J.Y., King, H.O., and Palmiter, R.D. (2021). Satb2 neurons in the parabrachial nucleus mediate taste perception. Nat. Commun. 12, 224. 10.1038/s41467-020-20100-8.

42. Ferreira, G., Ferry, B., Meurisse, M., and Lévy, F. (2006). Forebrain structures specifically activated by conditioned taste aversion. Behav. Neurosci. 120, 952–962. 10.1037/0735-7044.120.4.952.

43. Essner, R.A., Ruda, K., Choh, H.J., Kucukdereli, H., Amsalem, O., Edelhaus, J., Grødem, S., Lensjø, K.K., Lever, T.E., and Andermann, M.L. (2025). Brainstem sensing of multiple body signals during food consumption. Preprint at bioRxiv, 10.1101/2025.04.28.651046.

44. Schiavo, G., Matteoli, M., and Montecucco, C. (2000). Neurotoxins affecting neuroexocytosis. Physiol. Rev. 80, 717–766. 10.1152/physrev.2000.80.2.717.

45. Fischer, K.B., Collins, H.K., and Callaway, E.M. (2019). Sources of off-target expression from recombinase-dependent AAV vectors and mitigation with cross-over insensitive ATG-out vectors. Proc. Natl. Acad. Sci. U. S. A. 116, 27001–27010. 10.1073/pnas.1915974116.

46. Cheng, W., Gordian, D., Ludwig, M.Q., Pers, T.H., Seeley, R.J., and Myers, M.G. (2022). Hindbrain circuits in the control of eating behaviour and energy balance. Nat. Metab. 4, 826–835. 10.1038/s42255-022-00606-9.

47. Ramirez, I. (1984). Behavioral and physiological consequences of intragastric oil feeding in rats. Physiol. Behav. 33, 421–426. 10.1016/0031-9384(84)90164-1.

48. Corey, D.T. (1978). The determinants of exploration and neophobia. Neurosci. Biobehav. Rev. 2, 235–253. 10.1016/0149-7634(78)90033-7.

49. Revusky, S.H., and Bedarf, E.W. (1967). Association of Illness with Prior Ingestion of Novel Foods. Science 155, 219–220. 10.1126/science.155.3759.219.

50. De la Casa, G., and Lubow, R.E. (1995). Latent inhibition in conditioned taste aversion: the roles of stimulus frequency and duration and the amount of fluid ingested during preexposure. Neurobiol. Learn. Mem. 64, 125–132. 10.1006/nlme.1995.1051.

51. Bowen, A.J., Chen, J.Y., Huang, Y.W., Baertsch, N.A., Park, S., and Palmiter, R.D. (2020). Dissociable control of unconditioned responses and associative fear learning by parabrachial CGRP neurons. eLife *S*, e59799. 10.7554/eLife.59799.

52. Nardone, S., De Luca, R., Zito, A., Klymko, N., Nicoloutsopoulos, D., Amsalem, O., Brannigan, C., Resch, J.M., Jacobs, C.L., Pant, D., et al. (2024). A spatially-resolved transcriptional atlas of the murine dorsal pons at single-cell resolution. Nat. Commun. 15, 1966. 10.1038/s41467-024-45907-7.

53. Pauli, J.L., Chen, J.Y., Basiri, M.L., Park, S., Carter, M.E., Sanz, E., McKnight, G.S., Stuber, G.D., and Palmiter, R.D. (2022). Molecular and anatomical characterization of parabrachial neurons and their axonal projections. eLife 11, e81868. 10.7554/eLife.81868.

54. Herbert, H., Moga, M.M., and Saper, C.B. (1990). Connections of the parabrachial nucleus with the nucleus of the solitary tract and the medullary reticular formation in the rat. J. Comp. Neurol. 293, 540–580. 10.1002/cne.902930404.

55. Moga, M.M., Herbert, H., Hurley, K.M., Yasui, Y., Gray, T.S., and Saper, C.B. (1990). Organization of cortical, basal forebrain, and hypothalamic afferents to the parabrachial nucleus in the rat. J. Comp. Neurol. 295, 624–661. 10.1002/cne.902950408.

56. Kim, D.-I., Kang, S.J., Jhang, J., Jo, Y.S., Park, S., Ye, M., Pyeon, G.H., Im, G.-H., Kim, S.-G., and Han, S. (2024). Encoding opposing valences through frequency-dependent transmitter switching in single peptidergic neurons. Preprint at bioRxiv, 10.1101/2024.11.09.622790.

57. Yates, S.C., Groeneboom, N.E., Coello, C., Lichtenthaler, S.F., Kuhn, P.-H., Demuth, H.-U., Hartlage-Rübsamen, M., Roßner, S., Leergaard, T., Kreshuk, A., et al. (2019). QUINT: Workflow for Quantification and Spatial Analysis of Features in Histological Images From Rodent Brain. Front. Neuroinformatics 13. 10.3389/fninf.2019.00075.

58. Matikainen-Ankney, B.A., Earnest, T., Ali, M., Casey, E., Wang, J.G., Sutton, A.K., Legaria, A.A., Barclay, K.M., Murdaugh, L.B., Norris, M.R., et al. (2021). An open-source device for measuring food intake and operant behavior in rodent home-cages. eLife 10, e66173. 10.7554/eLife.66173.

59. Pnevmatikakis, E.A., and Giovannucci, A. (2017). NoRMCorre: An online algorithm for piecewise rigid motion correction of calcium imaging data. J. Neurosci. Methods 291, 83–94. 10.1016/j.jneumeth.2017.07.031.

