## Supplemental Statistics Table for "Ingestion-activated CGRP neurons control learning but not satiety"

Supplemental Table 1. Statistical Information for all Main Figures and Extended Data Figures.

| **Figure 1** | **Sample Size** | **Test** | **Statistics/p value** | **Posthoc comparisons** | **p value** |
|---|---|---|---|---|---|
| Figure 1f | n = 425 | Simple linear regression | R2 = 0.1304, F(1, 423) = 63.45, p < 0.0001 | | |
| Figure 1g | n = 55 | Two-tailed welch's t-test | t(55.31) = 7.172, p < 0.0001 | | |
| Figure 1h | Air n = 121 | Kolmogorov-Smirnov Test (2-sample) | p = 0.0003, h = 1 | | |
| | Ensure n = 242 | | | | |
| Figure 1j | sucralose n = 37 | Kruskal-Wallis | W =10.3, p =0.0357 | Saline vs. Sucralose | 0.0685 |
| | saline n = 23 | | | Saline vs. Ensure | >0.9999 |
| | Ensure n = 55 | | | Saline vs. Intralipid | 0.0754 |
| | Intralipid n = 34 | | | Saline vs. Glucose | >0.9999 |
| | glucose n = 20 | | | | |
| Figure 1k - activated | Ensure n = 242 | Fisher's Exact Test | | | |
| | Air n = 121 | Ensure vs. Air | 8.06E-08 | Bonferonni (15) | 1.21E-06 |
| | Saline n = 124 | Ensure vs. saline | 0.418873478 | Bonferonni (15) | 1 |
| | Sucralose n = 127 | Ensure vs. sucralose | 0.205368513 | Bonferonni (15) | 1 |
| | Glucose n = 77 | Ensure vs. glucose | 0.541688092 | Bonferonni (15) | 1 |
| | Intralipid n = 135 | Ensure vs. Intralipid | 0.319414177 | Bonferonni (15) | 1 |
| | | Air vs. saline | 3.79E-05 | Bonferonni (15) | 0.00056888 |
| | | Air vs. sucralose | 1.94E-09 | Bonferonni (15) | 2.91E-08 |
| | | Air vs. glucose | 6.66E-07 | Bonferonni (15) | 1.00E-05 |
| | | Air vs. Intralipid | 7.12E-09 | Bonferonni (15) | 1.07E-07 |
| | | Saline vs. sucralose | 0.055167507 | Bonferonni (15) | 0.8275126 |
| | | Saline vs. glucose | 0.220707017 | Bonferonni (15) | 1 |
| | | Saline vs. Intralipid | 0.105574563 | Bonferonni (15) | 1 |
| | | Sucralose vs. glucose | 0.7477405 | Bonferonni (15) | 1 |
| | | sucralose vs. Intralipid | 0.784797922 | Bonferonni (15) | 1 |
| | | Glucose vs. Intralipid | 0.873002949 | Bonferonni (15) | 1 |
| Figure 1k - inhibited | Air n = 121 | Ensure vs. Air | 0.138181127 | Bonferonni (15) | 1 |
| | Saline n = 124 | Ensure vs. saline | 0.017738888 | Bonferonni (15) | 0.26608332 |
| | Sucralose n = 127 | Ensure vs. sucralose | 0.145975203 | Bonferonni (15) | 1 |
| | Glucose n = 77 | Ensure vs. glucose | 0.721662552 | Bonferonni (15) | 1 |
| | Intralipid n = 135 | Ensure vs. Intralipid | 0.279819065 | Bonferonni (15) | 1 |
| | | Air vs. saline | 0.481602023 | Bonferonni (15) | 1 |
| | | Air vs. sucralose | 1 | Bonferonni (15) | 1 |
| | | Air vs. glucose | 0.119973113 | Bonferonni (15) | 1 |
| | | Air vs. Intralipid | 0.680624677 | Bonferonni (15) | 1 |
| | | Saline vs. sucralose | 0.485987427 | Bonferonni (15) | 1 |
| | | Saline vs. glucose | 0.030652889 | Bonferonni (15) | 0.45979334 |
| | | Saline vs. Intralipid | 0.199099781 | Bonferonni (15) | 1 |
| | | Sucralose vs. glucose | 0.127836237 | Bonferonni (15) | 1 |
| | | sucralose vs. Intralipid | 0.689284208 | Bonferonni (15) | 1 |
| | | Glucose vs. Intralipid | 0.291716598 | Bonferonni (15) | 1 |
| Figure 1l--activated | sucralose n = 37 | Kruskal-Wallis | W = 1.470, p = 0.6893 | | |
| | saline n = 23 | | | | |
| | Ensure n = 55 | | | | |
| | Intralipid n = 37 | | | | |
| | glucose n = 20 | | | | |
| Figure 1l--inhibited | Ensure n = 21 | Kruskal-Wallis | W = 1.632, p = 0.8974 | | |
| | Glucose n - 4 | | | | |
| | Intralipid n = 10 | | | | |
| | Sucralose n = 8 | | | | |
| | Saline n = 4 | | | | |
| | Empty n = 4 | | | | |
| Figure 1m | sucralose n = 37 | Kruskal-Wallis | H(4) = 0.3349, p = 0.9874 | | |
| | saline n = 23 | | | | |
| | Ensure n = 55 | | | | |
| | Intralipid n = 37 | | | | |
| | glucose n = 20 | | | | |

| **Figure 2** | **Sample Size** | **Test** | **Statistics/p value** | **Posthoc comparisons** | **p value** |
|---|---|---|---|---|---|
| Figure 2d | n = 20 | Two-tailed paired t-test | t(19) = 4.579, p =0.0002 | | |
| Figure 2e | n = 20 | Two-tailed paired t-test | t(19) = 2.967,p =0.0143 | | |
| Figure 2g | chow n = 6 | Two-tailed unpaired t-test | t(11) = 3.362, p = 0.0063 | | |
| | HFD n = 7 | | | | |
| Figure 2i | chow n = 210, HFD n = 204 | Exact Fisher Test (activated) | p = 0.5740 | bonferroni (2 comparisons) | |
| | | Exact Fisher Test (inhibited) | p = 0.0118 | bonferroni (2 comparisons) | |
| Figure 2k | chow n = 51 | Mann-Whitney (two-tailed) | U = 1303, p = 0.5329 | | |
| | HFD n =55 | | | | |

| | | | | | |
|---|---|---|---|---|---|
| Figure 2l | chow n = 51 | Mann-Whitney (two-tailed) | U = 1315, p = 0.5780 | | |
| | HFD n =55 | | | | |
| Figure 2m | n = 51 | Two-tailed welch's t-test | t(50.44) = 5.736, p <0.0001 | | |
| Figure 2n | chow n = 51 | Repeated-measure two-way ANOVA | | Sidak's Multiple Comparisons | |
| | HFD n = 55 | Food x Bout | F (1, 104) = 0.8493, p =0.3589 | Chow, bout vs. interbout | <0.0001 **** |
| | | Food | F (1, 104) = 0.8173, p =0.3681 | HFD, bouts vs. interbout | <0.0001 **** |
| | | Bout | F (1, 104) = 56.45, p <0.0001 | Bouts, chow vs. HFD | 0.447 |
| | | | | Interbout, chow vs. HFD | 0.8044 |
| Figure 2p, rapid | chow n = 51 | Repeated-measure two-way ANOVA | | Sidak's Multiple Comparisons | |
| | HFD n = 55 | Food x Time | F (1, 103) = 2.845, p =0.0947 | Chow, start vs. end | 0.0001 *** |
| | | Food | F (1, 103) = 0.1339, p =0.7152 | HFD, start vs. end | <0.0001 **** |
| | | Time | F (1, 103) = 58.34, p <0.0001 | Start, chow vs. HFD | 0.7917 |
| | | | | End, chow vs. HFD | 0.3959 |
| Figure 2p, delayed | chow, n = 15 | Repeated-measure two-way ANOVA | | Sidak's Multiple Comparisons | |
| | HFD n = 14 | Food x Time | F (1, 27) = 0.01656, p =0.8985 | Chow, start vs. end | <0.0001 **** |
| | | Food | F (1, 27) = 1.981, p =0.1707 | HFD, start vs. end | <0.0001 **** |
| | | Time | F (1, 27) = 115.5, p <0.0001 | Start, chow vs. HFD | 0.5575 |
| | | | | End, chow vs. HFD | 0.4496 |

| **Figure 3** | **Sample Size** | **Test** | **Statistics/p value** | **Posthoc comparisons** | **p value** |
|---|---|---|---|---|---|
| Figure 3c - inhibited | Saline n = 192 | Fisher's exact test | | | |
| | 1 mL Ensure n = 184 | 1 mL Ensure vs. saline | 9.69E-05 | bonferonni (corrected for 15) | 0.0015 ** |
| | 1.5 mL Ensure n = 16 | 1 mL Ensure vs. 1.5 mL Ensure | 0.2252 | bonferonni (corrected for 15) | 1 |
| | Intralipid n = 163 | 1 mL Ensure vs. Intralipid | 0.374 | bonferonni (corrected for 15) | 1 |
| | protein n = 133 | 1 mL Ensure vs. protein | 0.0721 | bonferonni (corrected for 15) | 1 |
| | glucose n = 121 | 1 mL Ensure vs. glucoose | 0.3612 | bonferonni (corrected for 15) | 1 |
| | | 1 mL saline vs. 1.5 mL Ensure | 3.50E-07 | bonferonni (corrected for 15) | <0.0001 **** |
| | | 1 mL saline vs. Intralipid | 0.0042 | bonferonni (corrected for 15) | 0.0637 |
| | | 1 mL saline vs. protein | 0.114 | bonferonni (corrected for 15) | 1 |
| | | 1 mL saline vs. glucose | 6.37E-07 | bonferonni (corrected for 15) | <0.0001 **** |
| | | 1.5 mL Ensure vs. Intralipid | 0.0309 | bonferonni (corrected for 15) | 0.4642 |
| | | 1.5 mL Ensure vs. protein | 0.0029 | bonferonni (corrected for 15) | 0.0438 * |
| | | 1.5 mL Ensure vs. glucose | 0.7972 | bonferonni (corrected for 15) | 1 |
| | | Intralipid vs. protein | 0.3686 | bonferonni (corrected for 15) | 1 |
| | | Intralipid vs. glucose | 0.0518 | bonferonni (corrected for 15) | 0.7777 |
| | | protein vs. glucose | 0.0061 | bonferonni (corrected for 15) | 0.0911 |
| Figure 3c - activated | Saline n = 192 | Fisher's exact test | | | |
| | 1 mL Ensure n = 184 | 1 mL Ensure vs. saline | 0.1762 | bonferonni (corrected for 15) | 1 |
| | 1.5 mL Ensure n = 16 | 1 mL Ensure vs. 1.5 mL Ensure | 0.0026 | bonferonni (corrected for 15) | 0.0391 * |
| | Intralipid n = 163 | 1 mL Ensure vs. Intralipid | 5.49E-09 | bonferonni (corrected for 15) | <0.0001 **** |
| | protein n = 133 | 1 mL Ensure vs. protein | 6.67E-06 | bonferonni (corrected for 15) | 0.0001 *** |
| | glucose n = 121 | 1 mL Ensure vs. glucose | 3.67E-08 | bonferonni (corrected for 15) | <0.0001 **** |
| | | 1 mL saline vs. 1.5 mL Ensure | 1.09E-05 | bonferonni (corrected for 15) | 0.0002 *** |
| | | 1 mL saline vs. Intralipid | 6.78E-13 | bonferonni (corrected for 15) | <0.0001 **** |
| | | 1 mL saline vs. protein | 5.03E-09 | bonferonni (corrected for 15) | <0.0001 **** |
| | | 1 mL saline vs. glucose | 3.04E-12 | bonferonni (corrected for 15) | <0.0001 **** |
| | | 1.5 mL Ensure vs. Intralipid | 0.0069 | bonferonni (corrected for 15) | 0.1029 |
| | | 1.5 mL Ensure vs. protein | 0.1087 | bonferonni (corrected for 15) | 1 |
| | | 1.5 mL Ensure vs. glucose | 0.023 | bonferonni (corrected for 15) | 0.3455 |
| | | Intralipid vs. protein | 0.324 | bonferonni (corrected for 15) | 1 |
| | | Intralipid vs. glucose | 0.654 | bonferonni (corrected for 15) | 1 |
| | | protein vs. glucose | 0.6278 | bonferonni (corrected for 15) | 1 |
| Figure 3d - activated | Saline n = 11 | Kruskal-Wallis Test | H(4) =21.98, p = 0.0002 | Dunn's Multiple Comparisons | |
| | Ensure n = 18 | | | Saline vs. 1 mL Ensure | >0.9999 |
| | Glucose n = 63 | | | Saline vs. Glucose | 0.1916 |
| | Protein n = 40 | | | Saline vs. Protein | 0.0046 ** |
| | Intralipid n = 58 | | | Saline vs. Intralipid | 0.0128 * |
| | | | | 1 mL Ensure vs. Glucose | 0.5621 |
| | | | | 1 mL Ensure vs. Protein | 0.0096 ** |
| | | | | 1 mL Ensure vs. Intralipid | 0.029 * |
| | | | | Glucose vs. Protein | 0.3477 |
| | | | | Glucose vs. Intralipid | >0.9999 |
| | | | | Protein vs. Intralipid | >0.9999 |
| Figure 3d - inhibited | Saline n = 9 | Kruskal-Wallis Test | H(4) =5.026, p =0.2846 | | |
| | Ensure n = 32 | | | | |
| | Glucose n = 41 | | | | |
| | Protein n = 13 | | | | |
| | Intralipid n = 22 | | | | |
| Figure 3f | Saline n = 11 | Kruskal-Wallis Test | H(4) = 12.79, p = 0.0123 | Dunn's Multiple Comparisons | |
| | Ensure n = 18 | | | Saline vs. 1 mL Ensure | >0.9999 |
| | Glucose n = 63 | | | Saline vs. Glucose | >0.9999 |
| | Protein n = 40 | | | Saline vs. Protein | 0.2029 |
| | Intralipid n = 58 | | | Saline vs. Intralipid | 0.2243 |
| | | | | 1 mL Ensure vs. Glucose | 0.7034 |
| | | | | 1 mL Ensure vs. Protein | 0.076 |
| | | | | 1 mL Ensure vs. Intralipid | 0.0777 |

| | | | | | |
|---|---|---|---|---|---|
| | | | | Glucose vs. Protein | >0.9999 |
| | | | | Glucose vs. Intralipid | >0.9999 |
| | | | | Protein vs. Intralipid | >0.9999 |
| Figure 3g | Saline n = 11 | Logrank (Mantel-Cox) test | $X_2(2)$ =1.169, p = 0.5574 | | |
| | Ensure n = 18 | | | | |
| | Ensure(1.5) n = 35 | | | | |
| Figure 3h - during IG | Ensure (1.0) n = 184 | Kolmogorov-smirnov test (two-sample) | p = 0.118, h = 0 | | |
| | Ensure (1.5) n = 161 | | | | |
| Figure 3h - post-IG | Ensure (1.0) n = 184 | Kolmogorov-smirnov test (two-sample) | p = 0.028, h = 1 | | |
| | Ensure (1.5) n = 161 | | | | |
| Figure 3j | Saline n = 7 | One-way ANOVA | F (3, 20) = 8.029, p = 0.0010 | Saline vs. Ensure (1mL) | 0.8424 |
| | Ensure (1.0) n = 4 | | | Saline vs. Ensure (1.5 mL) | 0.8424 |
| | Ensure (1.5) n = 7 | | | Saline vs. Intralipid | 0.0016 ** |
| | Intralipid n = 6 | | | | |

| Figure 4 | Sample Size | Test | Statistics/p value | Posthoc comparisons | p value |
|---|---|---|---|---|---|
| Figure 4c - Ensure | Ensure n = 28 | Kruskal-Wallis Test | H(3) =85.21, p <0.0001 | Dunn's Multiple Comparisons | |
| | LiCl n = 82 | | | None vs. Ensure | <0.0001 **** |
| | both n = 9 | | | None vs. LiCl | >0.9999 |
| | none n = 77 | | | None vs. Both | <0.0001 **** |
| Figure 4c - LiCl | Ensure n = 28 | Kruskal-Wallis Test | H(3) =146.1, p <0.0001 | Dunn's Multiple Comparisons | |
| | LiCl n = 82 | | | None vs. Ensure | >0.9999 |
| | both n = 9 | | | None vs. LiCl | <0.0001 **** |
| | none n = 77 | | | None vs. Both | <0.0001 **** |
| Figure 4f - Ensure | Ensure n = 10 | Kruskal-Wallis Test | H(3) =34.76, p <0.0001 | Dunn's Multiple Comparisons | |
| | DON n = 36 | | | None vs. Ensure | <0.0001 **** |
| | both n = 3 | | | None vs. DON | 0.6471 |
| | none n = 59 | | | None vs. Both | 0.0079 ** |
| Figure 4f - DON | Ensure n = 10 | Kruskal-Wallis Test | H(3) =71.94, p <0.0001 | Dunn's Multiple Comparisons | |
| | DON n = 38 | | | None vs. Ensure | >0.9999 |
| | both n = 3 | | | None vs. DON | <0.0001 **** |
| | none n = 59 | | | None vs. Both | 0.01 * |
| Figure 4h | lateral n = 5 | Two-tailed upaired t-test | t(10) = 3.206, p =0.0094 | | |
| | dorsal n = 7 | | | | |
| Figure 4i | Ensure | Fisher Exact Test | Fisher's exact test (two-sided) | p =0.0012 | |
| | LiCl | Fisher Exact Test | Fisher's exact test (two-sided) | p <0.0001 | |
| | IG Ensure | Fisher Exact Test | Fisher's exact test (two-sided) | p =0.5803 | |
| | Chow | Fisher Exact Test | Fisher's exact test (two-sided) | p = 0.6047 | |

| Figure 5 | Sample Size | Test | Statistics/p value | Posthoc comparisons | p value |
|---|---|---|---|---|---|
| Figure 5c - total licks | n = 7/group | Repeated Measure two-way ANOVA | | | |
| | | Virus x laser | F (1, 12) = 0.5232, p =0.4833 | | |
| | | Virus | F (1, 12) = 0.09001, p =0.7693 | | |
| | | Laser | F (1, 12) = 0.3567, p =0.5615 | | |
| Figure 5d -bout # | n = 7/group | Repeated Measure two-way ANOVA | | Sidak's multiple comparisons | |
| | | Virus x laser | F (1, 12) = 0.3534, p =0.5633 | mCherry, OFF vs. ON | 0.8357 |
| | | Virus | F (1, 12) = 1.231, p =0.2890 | GtACR, OFF vs. ON | 0.9488 |
| | | Laser | F (1, 12) = 0.03193, p =0.8612 | OFF, mCherry vs. GtACR | 0.3943 |
| | | | | ON, mCherry vs. GtACR | 0.6904 |
| Figure 5d -bout duration | n = 7/group | Repeated Measure two-way ANOVA | | Sidak's multiple comparisons | |
| | | Virus x laser | F (1, 12) = 1.016, p =0.3333 | mCherry, OFF vs. ON | 0.9597 |
| | | Virus | F (1, 12) = 1.044, p =0.3270 | GtACR, OFF vs. ON | 0.4618 |
| | | Laser | F (1, 12) = 0.4101, p =0.5339 | OFF, mCherry vs. GtACR | 0.8267 |
| | | | | ON, mCherry vs. GtACR | 0.3531 |
| Figure 5d -meal # | n = 7/group | Repeated Measure two-way ANOVA | | Sidak's multiple comparisons | |
| | | Virus x laser | F (1, 12) = 0.3980, p =0.5399 | mCherry, OFF vs. ON | 0.7778 |
| | | Virus | F (1, 12) = 1.108, p =0.3133 | GtACR, OFF vs. ON | 0.9646 |
| | | Laser | F (1, 12) = 0.08224, p =0.7792 | OFF, mCherry vs. GtACR | 0.778 |
| | | | | ON, mCherry vs. GtACR | 0.4109 |
| Figure 5d -meal duration | n = 7/group | Repeated Measure two-way ANOVA | | Sidak's multiple comparisons | |
| | | Virus x laser | F (1, 12) = 0.3450, p =0.5678 | mCherry, OFF vs. ON | 0.9834 |
| | | Virus | F (1, 12) = 0.4459, p =0.5169 | GtACR, OFF vs. ON | 0.7683 |
| | | Laser | F (1, 12) = 0.1246, p =0.7303 | OFF, mCherry vs. GtACR | 0.9998 |
| | | | | ON, mCherry vs. GtACR | 0.6221 |
| Figure 5e, total licks | n = 7/group | Repeated Measure two-way ANOVA | | | |
| | | Virus x laser | F (1, 12) = 3.391, p =0.0904 | | |
| | | Virus | F (1, 12) = 0.8463, p =0.3757 | | |
| | | Laser | F (1, 12) = 0.05556, p =0.8176 | | |
| Figure 5f - bout # | n = 7/group | Repeated Measure two-way ANOVA | | Sidak's multiple comparisons | |

| | | | | | |
|---|---|---|---|---|---|
| | | Virus x laser | F (1, 12) = 2.276, p =0.1573 | mCherry, OFF vs. ON | 0.882 |
| | | Virus | F (1, 12) = 0.7207, p =0.4125 | GtACR, OFF vs. ON | 0.0468 * |
| | | Laser | F (1, 12) = 4.638, p =0.0523 | OFF, mCherry vs. GtACR | 0.9108 |
| | | | | ON, mCherry vs. GtACR | 0.4021 |
| Figure 5f - bout duration | n = 7/group | Repeated Measure two-way ANOVA | | Sidak's multiple comparisons | |
| | | Virus x laser | F (1, 12) = 1.941, p =0.1889 | mCherry, OFF vs. ON | 0.0222 * |
| | | Virus | F (1, 12) = 0.4153, p =0.5314 | GtACR, OFF vs. ON | 0.5456 |
| | | Laser | F (1, 12) = 8.074, p =0.0149 | OFF, mCherry vs. GtACR | 0.4915 |
| | | | | ON, mCherry vs. GtACR | 0.9912 |
| Figure 5f - meal # | n = 7/group | Repeated Measure two-way ANOVA | | Sidak's multiple comparisons | |
| | | Virus x laser | F (1, 12) = 0.4139, p =0.5321 | mCherry, OFF vs. ON | 0.6686 |
| | | Virus | F (1, 12) = 1.048, p =0.3261 | GtACR, OFF vs. ON | 0.9958 |
| | | Laser | F (1, 12) = 0.2771, p =0.6082 | OFF, mCherry vs. GtACR | 0.4399 |
| | | | | ON, mCherry vs. GtACR | 0.9613 |
| Figure 5f - meal duration | n = 7/group | Repeated Measure two-way ANOVA | | Sidak's multiple comparisons | |
| | | Virus x laser | F (1, 12) = 0.04860, p =0.8292 | mCherry, OFF vs. ON | 0.7291 |
| | | Virus | F (1, 12) = 0.06027, p =0.8102 | GtACR, OFF vs. ON | 0.8997 |
| | | Laser | F (1, 12) = 0.6586, p =0.4329 | OFF, mCherry vs. GtACR | 0.9347 |
| | | | | ON, mCherry vs. GtACR | 0.9995 |
| Figure 5g | mCherry n = 6 | Repeated Measure two-way ANOVA | | Sidak's multiple comparisons | |
| | GtACR n = 5 | Virus x laser | F (1, 9) = 6.435, p =0.0319 | mCherry, OFF vs. ON | 0.9818 |
| | | Virus | F (1, 9) = 6.510, p =0.0311 | GtACR, OFF vs. ON | 0.0116 * |
| | | Laser | F (1, 9) = 7.686, p =0.0217 | | |
| Figure 5i | mCherry n = 9 | Repeated Measure two-way ANOVA | | First, mCherry vs. GtACR | 0.931 |
| | GtACR n = 10 | Test Day x Virus | F (1, 17) = 3.672, p =0.0723 | Second, mCherry vs. GtACR | 0.0448 * |
| | | Test Day | F (1, 17) = 26.92, p <0.0001 | mCherry, First vs. second day | 0.0003 *** |
| | | Virus | F (1, 17) = 2.602, p =0.1251 | GtACR, first vs. second day | 0.058 |
| Figure 5j | mCherry n = 9 | Mixed-effect two-way ANOVA | | Tukey's Multiple Comparisons | |
| | GtACR n = 10 | Day of treatment | F (2, 33) = 2.459, p = 1.011 | Day 1, mCherry vs. GtACR | 0.0133 * |
| | | Virus | F (1, 17) = 13.03, p = 0.0022 | Day 2, mCherry vs. GtACR | 0.0015 |
| | | Day of treatment x Virus | F (2, 33) = 1.136, p = 0.3335 | Day 3, mCherry vs. GtACR | 0.0004 |
| | | | | mCherry, Day 1 vs. 2 | 0.6796 |
| | | | | mCherry, Day 1 vs. 3 | >0.9999 |
| | | | | mCherry, Day 2 vs. 3 | 0.6734 |
| | | | | GtACR, Day 1 vs. 2 | 0.0771 |
| | | | | GtACR, Day 1 vs. 3 | 0.1036 |
| | | | | GtACR, Day 2 vs. 3 | 0.9877 |
| Figure 5k | mCherry n = 6 | Repeated Measure two-way ANOVA | | | |
| | GtACR n = 11 | Virus x laser | F (1, 15) = 0.002916, p =0.9576 | | |
| | | Virus | F (1, 15) = 0.08760, p =0.7713 | | |
| | | Laser | F (1, 15) = 2.578e-005, p =0.9960 | | |

| **Supplemental Figure 1** | **Sample Size** | **Test** | **Statistics/p value** | **Posthoc comparisons** | **p value** |
|---|---|---|---|---|---|
| Supplemental Figure 1c | Empty n = 5 | One-way ANOVA | F (5, 31) = 3.072, p = 0.0229 | Tukey's multiple comparisons test | |
| | Saline n = 5 | | | Empty Sipper vs. Saline | 0.7962 |
| | Ensure n = 10 | | | Empty Sipper vs. Ensure | 0.4047 |
| | Intralipid n = 7 | | | Empty Sipper vs. Intralipid | 0.9597 |
| | Glucose n = 4 | | | Empty Sipper vs. Glucose | >0.9999 |
| | Sucralose n = 7 | | | Empty Sipper vs. Sucralose | 0.6356 |
| | | | | Saline vs. Ensure | 0.021 * |
| | | | | Saline vs. Intralipid | 0.2844 |
| | | | | Saline vs. Glucose | 0.9113 |
| | | | | Saline vs. Sucralose | 0.0638 |
| | | | | Ensure vs. Intralipid | 0.8883 |
| | | | | Ensure vs. Glucose | 0.3632 |
| | | | | Ensure vs. Sucralose | 0.9996 |
| | | | | Intralipid vs. Glucose | 0.9214 |
| | | | | Intralipid vs. Sucralose | 0.9767 |
| | | | | Glucose vs. Sucralose | 0.5705 |
| Supplemental Figure 1e | n = 24 | Wilcoxon matched-pairs signed rank test | W = -42.0, p = 0.5646 | | |
| Supplemental Figure 1g | n = 55 | Two-tailed paired t-test | t(54) = 5.700, p < 0.0001 | | |
| Supplemental Figure 1h | caloric n = 112 | Repeated Measure Two-way ANOVA | | Sidak's Multiple comparisons | |
| | non-caloric n = 60 | Solution x Time | F (1, 170) = 1.721, p =0.1914 | Caloric, 0-10 vs. 20-30 | <0.0001 **** |
| | | Solution | F (1, 170) = 3.262, p=0.0727 | Non-caloric, 0-10 vs. 20-30 | 0.0002 *** |
| | | Time | F (1, 170) = 59.72, p <0.0001 | 0-10, Caloric vs. Non-caloric | 0.3889 |
| | | | | 20-30, Caloric vs. Non-caloric | 0.0634 |
| Supplemental Figure 1i | n = 150 | Wilcoxon matched-pairs signed rank test | W =789.0, p = 0.4609 | | |
| Supplemental Figure 1j | 0-1245, n =23 | Kruskal-Wallis | H(2) = 2.089, p = 0.3519 | Dunn's multiple comparisons test | |
| | 1246-2490, n = 51 | | | 0-1245 vs. 1246-2490 | 0.4492 |
| | 2492-3735, n = 41 | | | 0-1245 vs. 2492-3735 | >0.9999 |
| | | | | 1246-2490 vs. 2492-3735 | >0.9999 |

| Supplemental Figure 3 | Sample Size | Test | Statistics/p value | Posthoc comparisons | p value |
|---|---|---|---|---|---|
| Supplemental Figure 3b | chow n = 6 | Repeated Measure two-way ANOVA | | Sidak's Multiple Comparisons | |
| | HFD n = 7 | Time x Food | F (1, 11) = 2.893, p =0.1170 | Start, chow vs. HFD | 0.2818 |
| | | Time | F (1, 11) = 32.65, p =0.0001 | End, chow vs. HFD | 0.9358 |
| | | Food | F (1, 11) = 0.4650, p =0.5094 | Chow, start vs. end | 0.0007 *** |
| | | | | HFD, start vs. end | 0.0261 * |
| Supplemental Figure 3d, rapid | chow n = 13 | Repeated Measure two-way ANOVA | | | |
| | HFD n = 31 | Food x Time | F (1, 42) = 0.3605, p =0.5515 | | |
| | | Food | F (1, 42) = 0.3377, p =0.5643 | | |
| | | Time | F (1, 42) = 0.3197, p =0.5748 | | |
| Supplemental Figure 3d, delayed | chow n = 27 | Repeated Measure two-way ANOVA | | Sidak's Multiple Comparisons | |
| | HFD n = 24 | Food x Time | F (1, 49) = 4.907, p =0.0314 | Chow, early vs. late | <0.0001 **** |
| | | Food | F (1, 49) = 0.04869, p =0.8263 | HFD, early vs. late | <0.0001 **** |
| | | Time | F (1, 49) = 144.5, p <0.0001 | Early, chow vs. HFD | 0.196 |
| | | | | Late, chow vs. HFD | 0.346 |

| Supplemental Figure 4 | Sample Size | Test | Statistics/p value | Posthoc comparisons | p value |
|---|---|---|---|---|---|
| Supplemental Figure 4b | Glucose n = 63 | Logrank (Mantel-Cox) test | $X_2(2)$ = .542, p = 0.1701 | | |
| | Intralipid n = 58 | | | | |
| | Protein n = 40 | | | | |
| Supplemental Figure 4c | Glucose n = 63 | Repeated Measure two-way ANOVA | | Sidak's multiple comparisons | |
| | Intralipid n = 58 | Time x Solution | F (12, 948) = 10.93, p <0.0001 | -5-0: Glucose vs. Protein | 0.9991 |
| | Protein n = 40 | Tme | F (6, 948) = 51.54, p <0.0001 | -5-0: Glucose vs. Intralipid | 0.9987 |
| | | Solution | F (2, 158) = 3.107, p =0.0475 | -5-0: Protein vs. Intralipid | >0.9999 |
| | | | | 0-5: Glucose vs. Protein | 0.1084 |
| | | | | 0-5: Glucose vs. Intralipid | 0.1592 |
| | | | | 0-5: Protein vs. Intralipid | 0.9785 |
| | | | | 5-10: Glucose vs. Protein | 0.3793 |
| | | | | 5-10: Glucose vs. Intralipid | 0.4132 |
| | | | | 5-10: Protein vs. Intralipid | 0.9965 |
| | | | | 10-15: Glucose vs. Protein | <0.0001 **** |
| | | | | 10-15: Glucose vs. Intralipid | 0.9417 |
| | | | | 10-15: Protein vs. Intralipid | <0.0001 **** |
| | | | | 15-20: Glucose vs. Protein | <0.0001 **** |
| | | | | 15-20: Glucose vs. Intralipid | 0.3046 |
| | | | | 15-20: Protein vs. Intralipid | 0.0001 *** |
| | | | | 20-25: Glucose vs. Protein | 0.0103 * |
| | | | | 20-25: Glucose vs. Intralipid | 0.9675 |
| | | | | 20-25: Protein vs. Intralipid | 0.0352 * |
| | | | | 25-30: Glucose vs. Protein | 0.7852 |
| | | | | 25-30: Glucose vs. Intralipid | 0.7356 |
| | | | | 25-30: Protein vs. Intralipid | 0.2741 |
| Supplemental Figure 4d | Saline n = 9 | Kruskal-Wallis Test | H(4) = 1.127, p =0.8899 | | |
| | Ensure n = 32 | | | | |
| | Glucose n = 41 | | | | |
| | Protein n = 13 | | | | |
| | Intralipid n = 22 | | | | |
| Supplemental Figure 4e | Saline n = 9 | Kruskal-Wallis Test | H(4) = 5.221, p = 0.2654 | | |
| | Ensure n = 32 | | | | |
| | Glucose n = 41 | | | | |
| | Protein n = 13 | | | | |
| | Intralipid n = 22 | | | | |

| Supplemental Figure 5 | Sample Size | Test | Statistics/p value | Posthoc comparisons | p value |
|---|---|---|---|---|---|
| Supplemental Figure 5c | CCK10 n = 39 | Repeated Measure Two-way ANOVA | | Sidak's Multiple Comparisons | |
| | CCK30 n = 40 | Time x Dose | F (1.442, 111.0) = 0.2546, p =0.7015 | 10 vs 30: 600 | 0.2303 |
| | | Time | F (1.442, 111.0) = 7.166, p =0.0035 | 10 vs 30: 1200 | 0.0313 * |
| | | Dose | F (1, 77) = 7.327, p =0.0084 | 10 vs 30: 1800 | 0.0404 * |
| Supplemental Figure 5d | Ex4-3 n = 7 | Repeated Measure Two-way ANOVA | | Sidak's Multiple Comparisons | |
| | Ex4-150 n = 20 | Time x Dose | F (2, 50) = 0.6077, p =0.5486 | 3 vs 150: 600 | 0.465 |
| | | Time | F (2, 50) = 10.52, p =0.0002 | 3 vs 150: 1200 | 0.4595 |
| | | Dose | F (1, 25) = 1.551, p =0.2245 | 3 vs 150: 1800 | 0.6914 |

| Supplemental Figure 6 | Sample Size | Test | Statistics/p value | Posthoc comparisons | p value |
|---|---|---|---|---|---|
| Supplemental Figure 6b | n = 3 | Repeated Measure Two-way ANOVA | | Tukey's Multiple Comparison | |
| | | Gut area x Solution | F (2.985, 10.45) = 17.04, p =0.0002 | Stomach: Saline vs. Ensure | 0.0145 * |
| | | Gut area | F (1.492, 10.45) = 26.15, p =0.0002 | Stomach: Saline vs. Intralipid | 0.0349 * |
| | | Solution | F (2, 7) = 5.542, p =0.0361 | Stomach: Ensure vs. Intralipid | 0.2082 |
| | | | | 0-10 cm: Saline vs. Ensure | 0.1205 |
| | | | | 0-10 cm: Saline vs. Intralipid | 0.6938 |
| | | | | 0-10 cm: Ensure vs. Intralipid | 0.4173 |
| | | | | 11-20 cm: Saline vs. Ensure | 0.1432 |
| | | | | 11-20 cm: Saline vs. Intralipid | 0.1712 |
| | | | | 11-20 cm: Ensure vs. Intralipid | 0.9231 |
| Supplemental Figure 6c | n = 3 | Repeated Measure Two-way ANOVA | | Tukey's Multiple Comparison | |
| | | Gut area x Solution | F (4, 14) = 0.6922, p =0.6095 | Stomach: Saline vs. Ensure | 0.9878 |
| | | Gut area | F (2, 14) = 10.27, p =0.0018 | Stomach: Saline vs. Intralipid | 0.7411 |

| | | | | | |
|---|---|---|---|---|---|
| | | Solution | F (2, 7) = 0.6031, p =0.5732 | Stomach: Ensure vs. Intralipid | 0.8306 |
| | | | | 0-10 cm: Saline vs. Ensure | 0.9937 |
| | | | | 0-10 cm: Saline vs. Intralipid | 0.3735 |
| | | | | 0-10 cm: Ensure vs. Intralipid | 0.3177 |
| | | | | 11-20 cm: Saline vs. Ensure | 0.8347 |
| | | | | 11-20 cm: Saline vs. Intralipid | 0.8682 |
| | | | | 11-20 cm: Ensure vs. Intralipid | 0.9938 |
| Supplemental Figure 6d | n = 3 | One-way ANOVA | F (2, 7) = 20.86, p = 0.0011 | Tukey's Multiple Comparison | |
| | | | | Saline vs. Ensure | 0.001 ** |
| | | | | Saline vs. Lipid | 0.0058 ** |
| | | | | Ensure vs. Lipid | 0.1667 |
| Supplemental Figure 6e | n = 3 | One-way ANOVA | F (2, 7) = 12.25, p =0.0052 | Tukey's Multiple Comparison | |
| | | | | Saline vs. Ensure | 0.0058 ** |
| | | | | Saline vs. Lipid | 0.0139 * |
| | | | | Ensure vs. Lipid | 0.5717 |

| **Supplemental Figure 7** | **Sample Size** | **Test** | **Statistics/p value** | **Posthoc comparisons** | **p value** |
|---|---|---|---|---|---|
| Supplemental Figure 7a -total licks | mCherry n = 8 | Repeated measure two-way ANOVA | | | |
| | GtACR n = 7 | Virus x Drug | F (1, 13) = 0.6736, p =0.4266 | | |
| | | Main effect of virus | F (1, 13) = 0.2609, p =0.6181 | | |
| | | Main effect of drug | F (1, 13) = 0.0001038, p =0.9920 | | |
| Supplemental Figure 7b -Bout # | mCherry n = 8 | Repeated measure two-way ANOVA | | | |
| | GtACR n = 7 | Virus x Drug | F (1, 13) = 0.1070, p =0.7488 | | |
| | | Main effect of virus | F (1, 13) = 0.8038, p =0.3862 | | |
| | | Main effect of drug | F (1, 13) = 1.571, p =0.2321 | | |
| Supplemental Figure 7b -Bout duration | mCherry n = 8 | Repeated measure two-way ANOVA | | | |
| | GtACR n = 7 | Virus x Drug | F (1, 13) = 3.462, p =0.0856 | | |
| | | Main effect of virus | F (1, 13) = 0.7092, p =0.4149 | | |
| | | Main effect of drug | F (1, 13) = 0.7599, p =0.3992 | | |
| Supplemental Figure 7b -Meal # | mCherry n = 8 | Repeated measure two-way ANOVA | | | |
| | GtACR n = 7 | Virus x Drug | F (1, 13) = 0.09292, p =0.7653 | | |
| | | Main effect of virus | F (1, 13) = 0.3237, p =0.5791 | | |
| | | Main effect of drug | F (1, 13) = 0.09292, p =0.7653 | | |
| Supplemental Figure 7b -Meal duration | mCherry n = 8 | Repeated measure two-way ANOVA | | | |
| | GtACR n = 7 | Virus x Drug | F (1, 13) = 0.02299, p =0.8818 | | |
| | | Main effect of virus | F (1, 13) = 2.261, p =0.1565 | | |
| | | Main effect of drug | F (1, 13) = 1.062, p =0.3216 | | |
| Supplemental Figure 7c -total licks | mCherry n = 8 | Repeated measure two-way ANOVA | | | |
| | GtACR n = 10 | Virus x Drug | F (1, 16) = 0.2711, p =0.6097 | | |
| | | Main effect of virus | F (1, 16) = 1.286, p =0.2735 | | |
| | | Main effect of drug | F (1, 16) = 0.6100, p =0.4462 | | |
| Supplemental Figure 7d -Bout # | mCherry n = 8 | Repeated measure two-way ANOVA | | | |
| | GtACR n = 10 | Virus x Drug | F (1, 16) = 0.3348, p =0.5709 | mCherry, OFF vs. ON | 0.4651 |
| | | Main effect of virus | F (1, 16) = 1.175, p =0.2943 | GtACR, OFF vs. ON | 0.0923 |
| | | Main effect of drug | F (1, 16) = 5.231, p =0.0362 | OFF, mCherry vs. GtACR | 0.6675 |
| | | | | ON, mCherry vs. GtACR | 0.4101 |
| Supplemental Figure 7d -Bout duration | mCherry n = 8 | Repeated measure two-way ANOVA | | | |
| | GtACR n = 10 | Virus x Drug | F (1, 16) = 0.8896, p =0.3596 | | |
| | | Main effect of virus | F (1, 16) = 0.1174, p =0.7363 | | |
| | | Main effect of drug | F (1, 16) = 2.181, p =0.1591 | | |
| Supplemental Figure 7d -Meal # | mCherry n = 8 | Repeated measure two-way ANOVA | | | |
| | GtACR n = 10 | Virus x Drug | F (1, 16) = 1.886, p =0.1886 | | |
| | | Main effect of virus | F (1, 16) = 2.411, p =0.1400 | | |
| | | Main effect of drug | F (1, 16) = 1.886, p =0.1886 | | |
| Supplemental Figure 7d -Meal Duration | mCherry n = 8 | Repeated measure two-way ANOVA | | | |
| | GtACR n = 10 | Virus x Drug | F (1, 16) = 1.346, p =0.2630 | | |
| | | Main effect of virus | F (1, 16) = 2.971, p =0.1040 | | |
| | | Main effect of drug | F (1, 16) = 0.1084, p =0.7463 | | |

| **Supplemental Figure 8** | **Sample Size** | **Test** | **Statistic/p value** | **Posthoc comparisons** | **p value** |
|---|---|---|---|---|---|
| Supplemental Figure 8b -total licks | mCherry n = 6 | Repeated measure two-way ANOVA | | | |
| | hM4D n = 8 | Virus x Drug | F (1, 12) = 8.112, p =0.0147 | mCherry, Saline vs. DCZ | 0.5373 |
| | | Main effect of virus | F (1, 12) = 0.02196, p =0.8847 | hM4, Saline vs. DCZ | 0.0166 * |
| | | Main effect of drug | F (1, 12) = 1.636, p =0.2250 | Saline, mCherry vs. hM4 | 0.4769 |
| | | | | DCZ, mCherry vs. hM4 | 0.6546 |
| Supplemental Figure 8c -Bout # | mCherry n = 6 | Repeated measure two-way ANOVA | | | |
| | hM4D n = 8 | Virus x Drug | F (1, 12) = 16.55, p =0.0016 | mCherry, Saline vs. DCZ | 0.0924 |
| | | Main effect of virus | F (1, 12) = 5.885e-005, p =0.9940 | hM4, Saline vs. DCZ | 0.0065 ** |
| | | Main effect of drug | F (1, 12) = 0.5296, p =0.4807 | Saline, mCherry vs. hM4 | 0.2845 |
| | | | | DCZ, mCherry vs. hM4 | 0.2911 |
| Supplemental Figure 8c -Bout duration | mCherry n = 6 | Repeated measure two-way ANOVA | | | |

| | | | |
|---|---|---|---|
| | hM4D n = 8 | Virus x Drug | F (1, 12) = 1.977, p =0.1851 |
| | | Main effect of virus | F (1, 12) = 0.2837, p =0.6040 |
| | | Main effect of drug | F (1, 12) = 0.9365, p =0.3523 |
| Supplemental Figure 8c -Meal # | mCherry n = 6 | Repeated measure two-way ANOVA | |
| | hM4D n = 8 | Virus x Drug | F (1, 12) = 0.02210, p =0.8843 |
| | | Main effect of virus | F (1, 12) = 0.01292, p =0.9114 |
| | | Main effect of drug | F (1, 12) = 1.415, p =0.2573 |
| Supplemental Figure 8c -Meal duration | mCherry n = 6 | Repeated measure two-way ANOVA | |
| | hM4D n = 8 | Virus x Drug | F (1, 12) = 0.7307, p =0.4094 |
| | | Main effect of virus | F (1, 12) = 0.07716, p =0.7859 |
| | | Main effect of drug | F (1, 12) = 2.203, p =0.1636 |
| Supplemental Figure 8d -total licks | mCherry n = 6 | Repeated measure two-way ANOVA | |
| | hM4D n = 8 | Virus x Drug | F (1, 12) = 0.1054, p =0.7511 |
| | | Main effect of virus | F (1, 12) = 0.003984, p =0.9507 |
| | | Main effect of drug | F (1, 12) = 0.01472, p =0.9054 |
| Supplemental Figure 8e -Bout # | mCherry n = 6 | Repeated measure two-way ANOVA | |
| | hM4D n = 8 | Virus x Drug | F (1, 12) = 1.238, p =0.2876 |
| | | Main effect of virus | F (1, 12) = 2.930, p =0.1126 |
| | | Main effect of drug | F (1, 12) = 2.216, p =0.1624 |
| Supplemental Figure 8e -Bout duration | mCherry n = 6 | Repeated measure two-way ANOVA | |
| | hM4D n = 8 | Virus x Drug | F (1, 12) = 0.0008501, p =0.9772 |
| | | Main effect of virus | F (1, 12) = 0.2043, p =0.6593 |
| | | Main effect of drug | F (1, 12) = 3.347, p =0.0923 |
| Supplemental Figure 8e -Meal # | mCherry n = 6 | Repeated measure two-way ANOVA | |
| | hM4D n = 8 | Virus x Drug | F (1, 12) = 1.421, p =0.2563 |
| | | Main effect of virus | F (1, 12) = 3.045, p =0.1065 |
| | | Main effect of drug | F (1, 12) = 1.115, p =0.3119 |
| Supplemental Figure 8e -Meal duration | mCherry n = 6 | Repeated measure two-way ANOVA | |
| | hM4D n = 8 | Virus x Drug | F (1, 12) = 0.3450, p =0.5678 |
| | | Main effect of virus | F (1, 12) = 0.02288, p =0.8823 |
| | | Main effect of drug | F (1, 12) = 1.836, p =0.2004 |
| Supplemental Figure 8f | mCherry n = 6 | Repeated measure two-way ANOVA | |
| | | Time | F (2, 10) = 110.0, p <0.0001 |
| | | Drug | F (1, 5) = 0.1138, p =0.7495 |
| | | Time x Drug | F (2, 10) = 1.069, p =0.3796 |
| | hM4D n = 7 | Repeated measure two-way ANOVA | |
| | | Time | F (2, 12) = 104.0, p <0.0001 |
| | | Drug | F (1, 6) = 1.115, p =0.3317 |
| | | Time x Drug | F (2, 12) = 0.1824, p =0.8355 |
| Supplemental Figure 8g | mCherry n = 6 | Repeated measure two-way ANOVA | |
| | hM4D n = 7 | Virus x Drug | F(1, 11) = 1.452, p = 0.2535 |
| | | Main effect of virus | F(1, 11) = 0.9165, p = 0.3590 |
| | | Main effect of drug | F(1, 11) = 0.06804, p = 0.7990 |
| Supplemental Figure 8h - total intake | mCherry n = 3 | Repeated measure two-way ANOVA | |
| | hM4D n = 5 | Virus x Drug | F (1, 6) = 0.01186, p =0.9168 |
| | | Main effect of virus | F (1, 6) = 2.087, p =0.1987 |
| | | Main effect of drug | F (1, 6) = 0.03712, p =0.8536 |
| Supplemental Figure 8h - Meal # | mCherry n = 3 | Repeated measure two-way ANOVA | |
| | hM4D n = 5 | Virus x Drug | F (1, 6) = 0.09698, p =0.7660 |
| | | Main effect of virus | F (1, 6) = 2.223, p =0.1866 |
| | | Main effect of drug | F (1, 6) = 0.8728, p =0.3862 |
| Supplemental Figure 8h - Pellets/meal | mCherry n = 3 | Repeated measure two-way ANOVA | |
| | hM4D n = 5 | Virus x Drug | F (1, 6) = 0.01403, p =0.9096 |
| | | Main effect of virus | F (1, 6) = 0.09727, p =0.7657 |
| | | Main effect of drug | F (1, 6) = 3.458, p =0.1123 |
| Supplemental Figure 8j | mCherry n = 6 | Unpaired t-test with Welch's correction | t(6.343) = 4.870, p = 0.0024 |
| | hM4D n = 6 | | |
